# Genomics-FM: Universal Foundation Model for Versatile and Data-Efficient Functional Genomic Analysis

**DOI:** 10.1101/2024.07.16.603653

**Authors:** Peng Ye, Weiqing Bai, Yuchen Ren, Wenran Li, Lifeng Qiao, Chaoqi Liang, Linxiao Wang, Yuchen Cai, Jianle Sun, Zejun Yang, Peng Zheng, Nanqing Dong, Tao Chen, Zhihui Wang, Xihui Liu, Xinzhu Ma, Hongliang Yan, Zhen Wang, Sijia Wang, Wanli Ouyang

## Abstract

Artificial intelligence (AI) plays a crucial role in functional genomic analysis, offering great potential for comprehending biological phenomena such as heredity, development, diseases, and evolution. However, the development of AI models needs substantial labeled data, and these models are typically task-specific with limited generalizability to various applications. Here, we develop Genomics-FM, a genomic vocabulary driven foundation model that enables versatile and label-efficient functional genomic analysis. Specifically, Genomics-FM is first pretrained with ensemble genomic vocabulary on vast unlabelled data to learn comprehensive and generalizable representations and then finetuned with specific genomic vocabulary on limited labeled data to selectively activate and adapt the pretraining knowledge for specific tasks. We show that Genomics-FM significantly reduces the dependence on labeled data, and demonstrates the capability to outperform existing models across a comprehensive suite of tasks including genome annotation, epigenomic and expression profile prediction, and variant effect assessment. Remarkably, Genomics-FM even shows impressive zero-shot predictive capabilities across diverse species and tissues and exhibits noticeable adaptability to RNA-related tasks. With feasibility in data scarcity and even cross-domain biological scenarios, Genomics-FM will promote the broad application of AI and empower researchers to tackle previously insurmountable challenges, paving the way for groundbreaking research and discoveries.

## 1 Introduction

Artificial Intelligence (AI) has catalyzed substantial advancements in functional genomic analysis, as evidenced by the development of sophisticated AI models (Wang et al., 2023). These models have enhanced our understanding in areas including disease mechanisms (Chen et al., 2024), drug discovery (Cappelluti et al., 2024), phenotype prediction (Rozowsky et al., 2023; Zhang et al., 2022b), and evolutionary biology (Foley et al., 2023; Green et al., 2010). Traditionally, these models have relied on extensive volumes of high-quality, labeled data, requiring meticulous expert assessment and significant labor. However, the limited availability of domain-specific experts, combined with the vast and complex nature of genomic data, often results in substantial amounts of significant genomic datasets remaining unlabeled and underutilized (Whalen et al., 2022). This situation presents both a challenge and an opportunity for innovation in genomic research.

Foundation models provide a promising approach to addressing the data inefficiency challenge in genomics (Ji et al., 2021; Avsec et al., 2021; Dalla-Torre et al., 2023; Fishman et al., 2023; Theodoris et al., 2023; Cui et al., 2024). These models are typically pretrained on a broad array of unlabeled data by developing supervisory signals intrinsically present within the data itself, enabling the development of general-purpose feature representations. Subsequently, these models can be easily finetuned for specific tasks using limited labeled data. This capability is particularly valuable in genomics, a field characterized by an abundance of unlabeled data, diverse tasks, and a scarcity of labeled samples (Grosse & Gudgeon, 2021; Gürsoy et al., 2022). While some works have shown that foundation models can increase performance for specific genomic tasks, they often fail to perform consistently well across different genomic tasks, and there has been limited work demonstrating their label efficiency. These problems are largely attributed to not fully considering the unique characteristics of genomic data and tasks to customize foundation models.

In this paper, we introduce Genomics-FM, a foundation model driven by genomic vocabulary tailored to enhance versatile and label-efficient functional genomic analysis. Genomic vocabulary, analogous to a lexicon in linguistics (Vu et al., 2023), defines the conversion of continuous genomic sequences into discrete units (tokens) for model input (Robert et al., 2022; Dotan et al., 2024), which directly influences the representation of genomic data and the performance of AI models. Distinct from previous works that typically employ a single (Lin et al., 2023) and specific genomic vocabulary (Ji et al., 2021; Zhou et al., 2023b; Dalla-Torre et al., 2023) for both pretraining and fine-tuning, Genomics-FM constructs an ensemble genomic vocabulary that includes multiple vocabularies during pretraining, and selectively activates specific genomic vocabularies for the fine-tuning of different tasks. Such a strategy not only enriches the general-purpose genomic knowledge extracted during pretraining, but also ensures superior task-specific adaptation during fine-tuning. Consequently, our approach enhances both the performance and label efficiency of Genomics-FM, enabling it to excel across a broad spectrum of genomic tasks and applications with minimal labeled data requirements.

The realm of genomics spans a wide array of intricate and diverse studies (Wong et al., 2021; Novakovsky et al., 2023), each involving unique genomic tasks with distinct characteristics and preferences. A natural advantage of our design is its ability to perform well on various genomic tasks. To comprehensively evaluate our model, we collect data from more than 16 genomic tasks, encompassing multiple levels of annotating genetic elements, predicting gene expression levels, and assessing the impact of genetic variants. We find that Genomics-FM achieves the best results across all of these tasks compared to existing AI models. Further, we apply it to genomic tasks in transcriptomics (Dahlberg, 1989; Nirenberg & Leder, 1964; Wong & Clayton, 1986; Roundtree et al., 2017), where it still demonstrates promising performance, even surpassing existing large-scale RNA models (Chen et al., 2022). These results indicate that Genomics-FM can perform well in various tasks previously dominated by different domain-specific models, underscoring its potential to promote the broad application of AI across various domains.

Across various genomic tasks, labeled data are exceptionally scarce, and for some tasks, labeled data may be entirely absent. Another natural advantage of our design is its ability to enhance label efficiency, achieving similar performance to other AI models with much less labeled data. Notably, we show that Genomics-FM enables cross-species and crosstissue prediction. Specifically, while whole-genome sequences have been identified for many species, many important properties of genomic sequences, such as regulatory elements classes, chromatin accessibility, and gene expression, have not been well annotated except for a few model organisms or tissues (Przybyla & Gilbert, 2022; Lucas & Novoa, 2023). We demonstrate that, with only human training data, our model can generalize well to multiple species, even surpassing the within-species training results of previous methods. Similarly, trained only on data from a limited set of tissues, it can effectively predict gene expression and regulatory elements in entirely different tissues with higher accuracy than traditional tissue-specific models. These applications collectively highlight the model’s adaptability in data scarcity and even cross-domain biological scenarios.

## 2 Results

### 2.1 Genomics-FM pretraining and finetuning

As shown in Fig. 1a, establishing a foundation model typically consists of two stages: unsupervised pretraining and supervised finetuning. At the unsupervised pretraining stage, the model is trained (pretrained) on vast unlabeled data with the Masked language modeling (MLM) (Kenton & Toutanova, 2019) objective. At the supervised finetuning stage, the model is further trained (finetuned) on limited labeled data with a specific supervised task. The primary difference between our Genomics-FM and previous DNA foundation models is the utilization of genomic vocabulary. Considering different genomic vocabularies generally capture distinct aspects of the genetic sequence, we construct an ensemble genomic vocabulary that encompasses multiple different vocabularies for unsupervised pretraining and adopts a specific genomic vocabulary suitable for the given downstream task during supervised fine-tuning.

**Figure 1.**
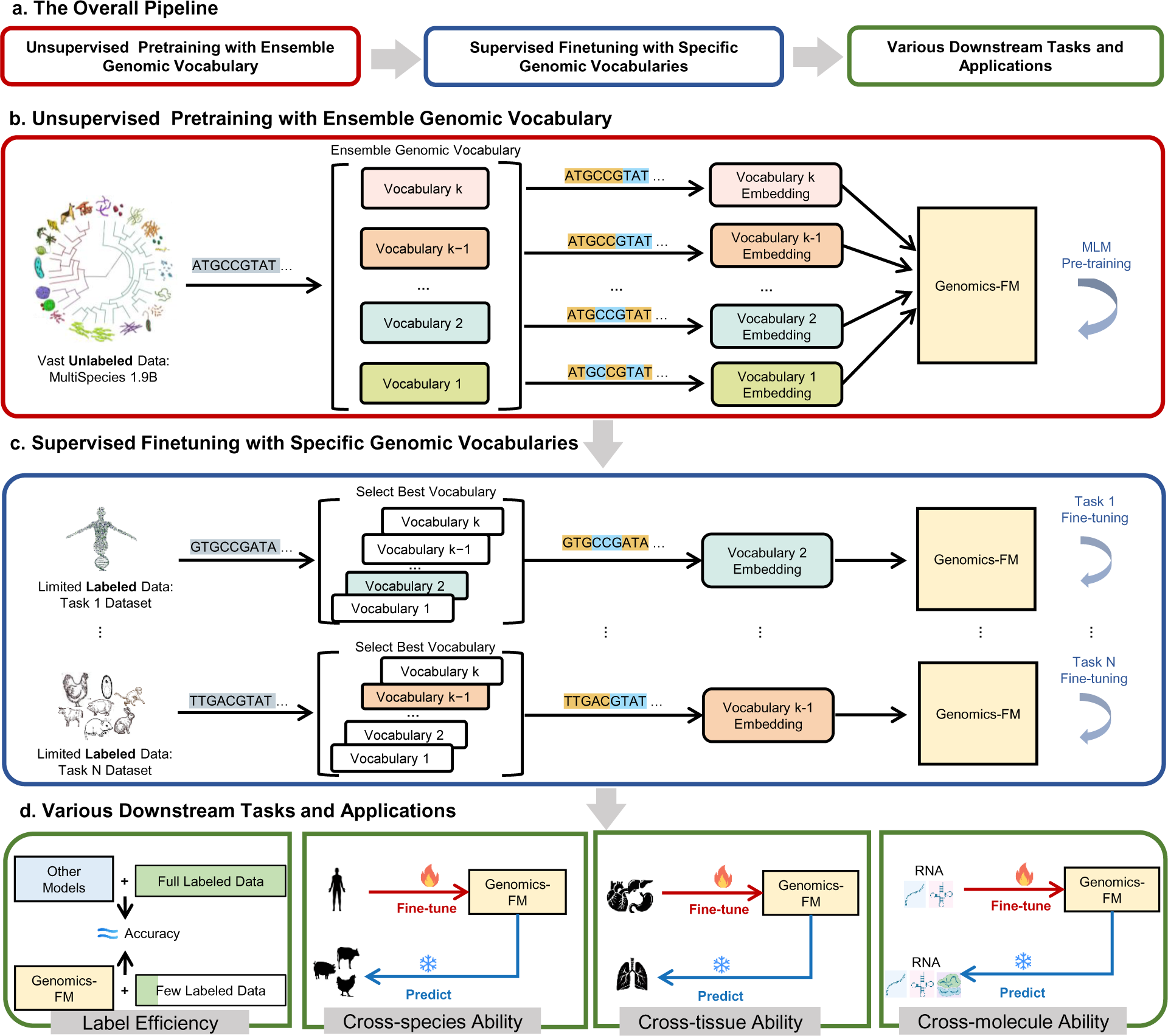
Schematic of development and applications of Genomics-FM. **a,** The overall pipeline of establishing a foundation model, including unsupervised pretraining on vast unlabeled data and supervised finetuning on limited labeled data. Here, we develop Genomics-FM, a genomic vocabulary driven foundation model, to enable versatile and label-efficient functional genomic analysis. The main difference between Genomics-FM and previous methods is the utilization of genomic vocabulary that defines how genetic sequences are transformed into a series of tokens for model input. **b,** During unsupervised pretraining, Genomics-FM employs a ensemble vocabulary that includes multiple different vocabularies to learn different types of genetic representations via masked language modeling. **c,** During supervised finetuing, Genomics-FM selectively activates the best vocabulary to extract the most suitable representation for specific downstream tasks. **d,** Highlight of Genomics-FM: 1) utilize much less labeled data to achieve performance comparable to previous state-of-the-art methods; demonstrate the capability for 2) cross-species analysis, 3) cross-tissue analysis, and 4) cross-molecule analysis.

#### Unsupervised pretraining with ensemble genomic vocabulary

During unsupervised pretraining, we construct an ensemble genomic vocabulary by integrating multiple different vocabularies, as shown in Fig. 1b. To enable joint training, the input embeddings of Genomics-FM are initialized with the ensemble vocabulary by removing duplicates from multiple vocabularies. To distinguish between different vocabularies during pretraining, each DNA sequence is appended with a special token (aka, prompt) related to the specific vocabulary, allowing the model to recognize and adapt to the specific vocabulary. A large-scale multi-species unlabelled DNA sequence dataset, comprising 1160 species and the human 1KG dataset (Byrska-Bishop et al., 2022), containing a total of 1.9 billion sequences, is collected as the pretraining dataset, see Supplementary for more details.

#### Supervised finetuning with specific genomic vocabulary

Since different downstream tasks in genomic sequence analysis often exhibit different characteristics, we selectively activate the most appropriate vocabulary for each downstream task during finetuning, as illustrated in Fig. 1c. In detail, we activate only the prompt token associated with the optimal vocabulary and equip the model with a task-specific head, allowing it to specialize and excel in the given task with a limited amount of labeled data. The designs described above not only facilitate a better understanding of both labeled and unlabeled genomic sequences but also enhance the model’s adaptability to handle diverse downstream genomic tasks.(Fig. 1d). Moreover, this approach is flexible and highly scalable, allowing for the seamless incorporation of new vocabularies into the existing collection.

### 2.2 Genomics-FM for versatile genomics tasks

We first thoroughly validated the performance of Genomics-FM by integrating 16 genomic tasks and their publicly available datasets, which encompass a broad spectrum across various scales including regulatory element annotation, gene expression and chromatin state, variations, and species-level analyses. The compared methods consist of the mainstream DNA foundation models, including DNABERT (Ji et al., 2021), DNABERT2 (Zhou et al., 2023b), Nucleotide Transformer (NT) (Dalla-Torre et al., 2023), and HyenaDNA (Nguyen et al., 2024).

#### State-of-the-art performance across 16 genomics tasks

As shown in Fig. 2a, Genomics-FM can achieve notable performance on 16 tasks across different biology levels with a large lead. For the genomic annotation tasks, in both promoter prediction and splices site recognition tasks, Genomics-FM outperforms the second place (HyenaDNA and DNABERT, respectively) by increasing MCC of 2.57% and 3.32%, respectively. For the sequence property prediction task, namely yeast epigenetic marks prediction, Genomics-FM also exhibits strong capacities and obtains an averaged improvement of 3.69% MCC compared to the second rank (DNABERT2). Impressively, Genomics-FM also showcases its capabilities in complex tasks where the efficacy of foundation models was previously unexplored. As depicted in Fig. 2b, while existing DNA foundation models perform worse than Expecto (Zhou et al., 2018) and DeepStarr (de Almeida et al., 2022) for gene expression prediction and regulator property prediction tasks respectively, Genomics-FM still exhibits remarkable improvements of 6.35% in *R*^2^ and 2.59% in PCC respectively. Overall, as shown by the averaged performance across all 16 tasks in Fig. 2c, Genomics-FM obtains an average performance of 79.49%, significantly outperforming DNABERT, DNABERT2, HyenaDNA, and NT (which achieve average scores of 72.32%, 73.72%, 72.97%, and 74.19%, respectively). This demonstrates the remarkable effectiveness across various genomic tasks delivered by Genomics-FM.

**Figure 2.**
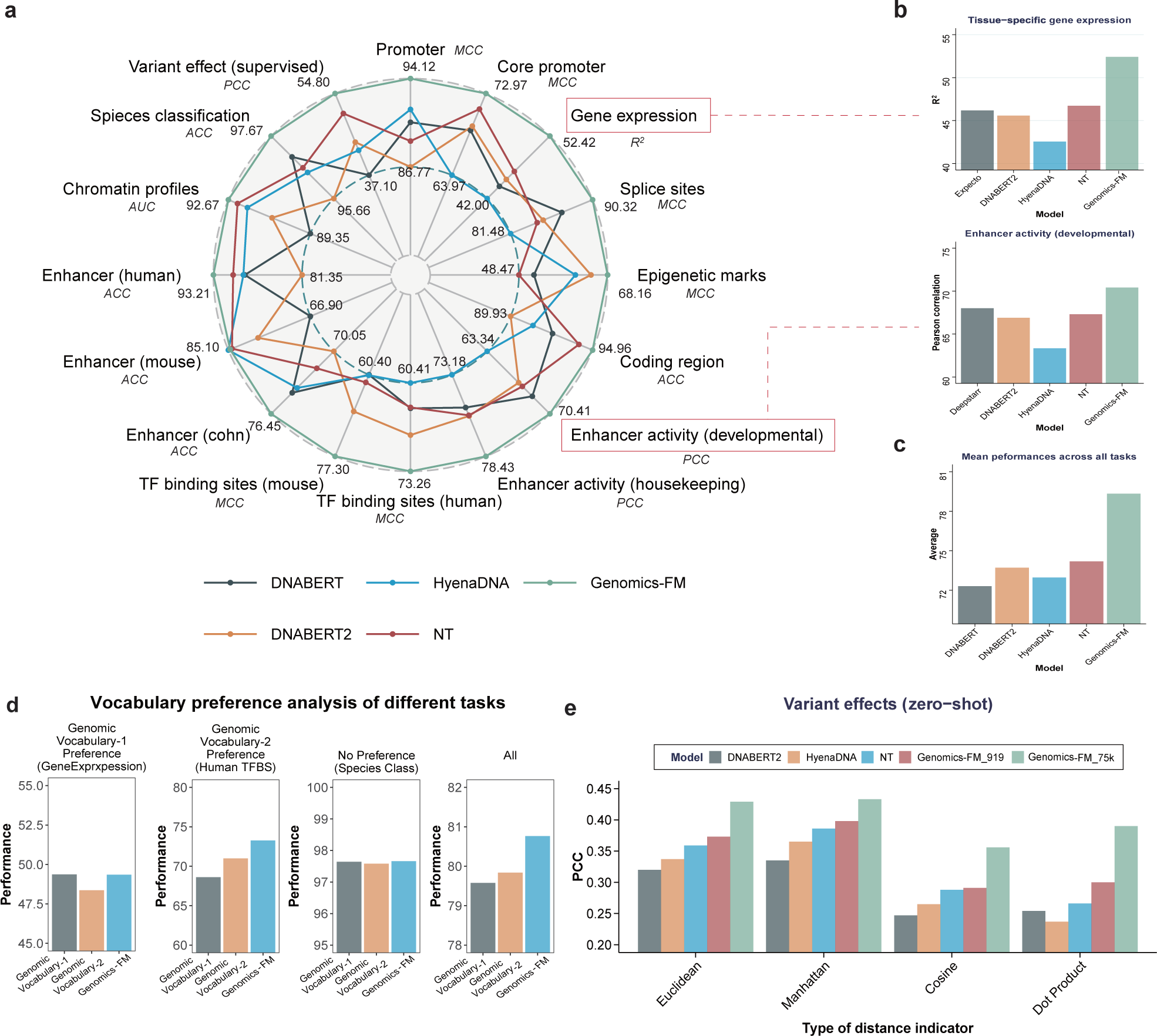
Evaluation of Genomics-FM on comprehensive genomic benchmark. **a,** Comprehensive evaluations across up to 16 genomic tasks, Genomics-FM consistently achieves the best performance. **b,** Detailed results on complex genomic tasks including developmental enhancer activity and cell-type specific gene expression prediction, Genomics-FM performs effectively on these tasks while other DNA foundation models falter. **c,** Mean performance across 16 tasks, Genomics-FM significantly surpasses other methods. **d,** Ablation studies verifying that different tasks prefer different vocabularies, Genomics-FM consistently performs well across all tasks. **e,** Generalization ability validation of Genomics-FM in zero-shot variant effect prediction, Genomics-FM can obtain better performance compared to other DNA foundation models.

#### The merits of the genomic vocabulary design in Genomics-FM

To show the effectiveness of the genomic vocabulary design of Genomics-FM, we compare three foundation models that differ only in their vocabulary. Genomic Vocabulary-1 and Genomic Vocabulary-2 refer to models pretrained and finetuined with a Byte-Pair Encoding vocabulary and K-mer (K=6) vocabulary respectively, both vocabularies are widely used in various pretrained models (Radford et al., 2018; 2019; Zhou et al., 2023b; Ji et al., 2021; Dalla-Torre et al., 2023). Genomics-FM, the proposed method, represents the model pretrained with an ensemble vocabulary and selectively finetuned with a specific vocabulary. As depicted in Fig. 2d, Genomic Vocabulary-2 generally excels in tasks requiring high-resolution information, such as transcription factor binding site (TFBS) identification, whereas Genomic Vocabulary-1 is more adept at tasks requiring whole-sequence information, e.g. estimating gene expression level. More similar results that verify the vocabulary preference across various tasks can be found in Supplementary. On the one hand, Genomics-FM can effectively solve the vocabulary preference problem by flexibly activating the vocabulary for fine-tuning, as verified by the average performance improvement of all tasks. On the other hand, Genomics-FM can learn more abundant general representation by pretraining with an ensemble vocabulary, as evidenced by comparable and even better performance when finetuned with the same vocabulary of Genomic Vocabulary-1 and Genomic Vocabulary-2.

#### Zero-shot variant effect prediction

We further compare the zero-shot variant effect prediction abilities of different methods. This experiment involves two steps: finetuning on epigenomic features, and zero-shot variant effect prediction. The pre-trained models are fine-tuned using two different sets of epigenetic features: 919 features from DeepSEA (Zhou & Troyanskaya, 2015) and more comprehensively 75,000 features collected from ChIP-Atlas (Zou et al., 2022). This results in two distinct models, Genomics-FM 919 and Genomics-FM 75k. During the zero-shot prediction stage, we use CAGI5 data (Shigaki et al., 2019; Dalla-Torre et al., 2023) to calculate the distance of predicted epigenomic profiles between sequences with different alleles. The correlation with effects measured by massively parallel reporter assays (MPRAs) experiments (Shigaki et al., 2019) is computed to evaluate zero-shot prediction performance. Four metrics (Euclidean, Manhattan, Cosine, and Dot product) are used, and the results are shown in Fig. 2e. Under the same settings, it is evident that Genomics-FM 919 consistently outperforms other DNA foundation models, demonstrating the superior zero-shot transfer ability of our model. Impressively, Genomics-FM 75k further achieves an astonishing performance of 0.429 and 0.433 when using Euclidean and Manhattan distance metrics respectively, surpassing many supervised learning approaches in the original competition (Shigaki et al., 2019). These findings underscore a robust correlation between the variant effects observed in the wet lab (ground truth) and those anticipated by Genomics-FM under a zero-shot scenario, highlighting the model’s capacity for generalization.

### 2.3 Genomics-FM substantially improves label efficiency

Label efficiency refers to the ability of AI models to achieve high performance with fewer labeled samples (annotations) during training, which saves time and money on wet-lab experiments. Better label efficiency can improve the applicability of AI models in data-limited applications and scenarios (Luo et al., 2017; Zhou et al., 2023a), which could raise great interest in biology fields (Gündüz et al., 2023; Theodoris et al., 2023) since lack of annotations is ubiquitous.

#### Superior Label efficiency of Genomics-FM

As depicted in Fig. 3a, Genomics-FM showcases comparable and even superior performance when trained with only 10% to 50% of the labeled data compared to previous best foundation models, including HyenaDNA, DNABERT, and Nucleotide Transformer (NT). Remarkably, in the Species classification dataset, 10% of labeled data (a 90% reduction in annotations) is sufficient to obtain competitive results. The average improvements in label efficiency across all tasks compared to representative DNA foundation models (DNA FMs) are also presented in Fig. 3b. Genomics-FM astonishingly reduces the need for annotations by 8.05, 4.65, and 5.01 times to achieve comparable performances with HyenaDNA, DNABERT2, and NT, respectively. While all these foundation models benefit from pretraining on vast unlabeled data, the proposed Genomics-FM, utilizing unified vocabulary for pretraining and selecting proper vocabulary for finetuning, exhibits the capability to more effectively make use of annotations and obtains a significant improvement in label efficiency.

**Figure 3.**
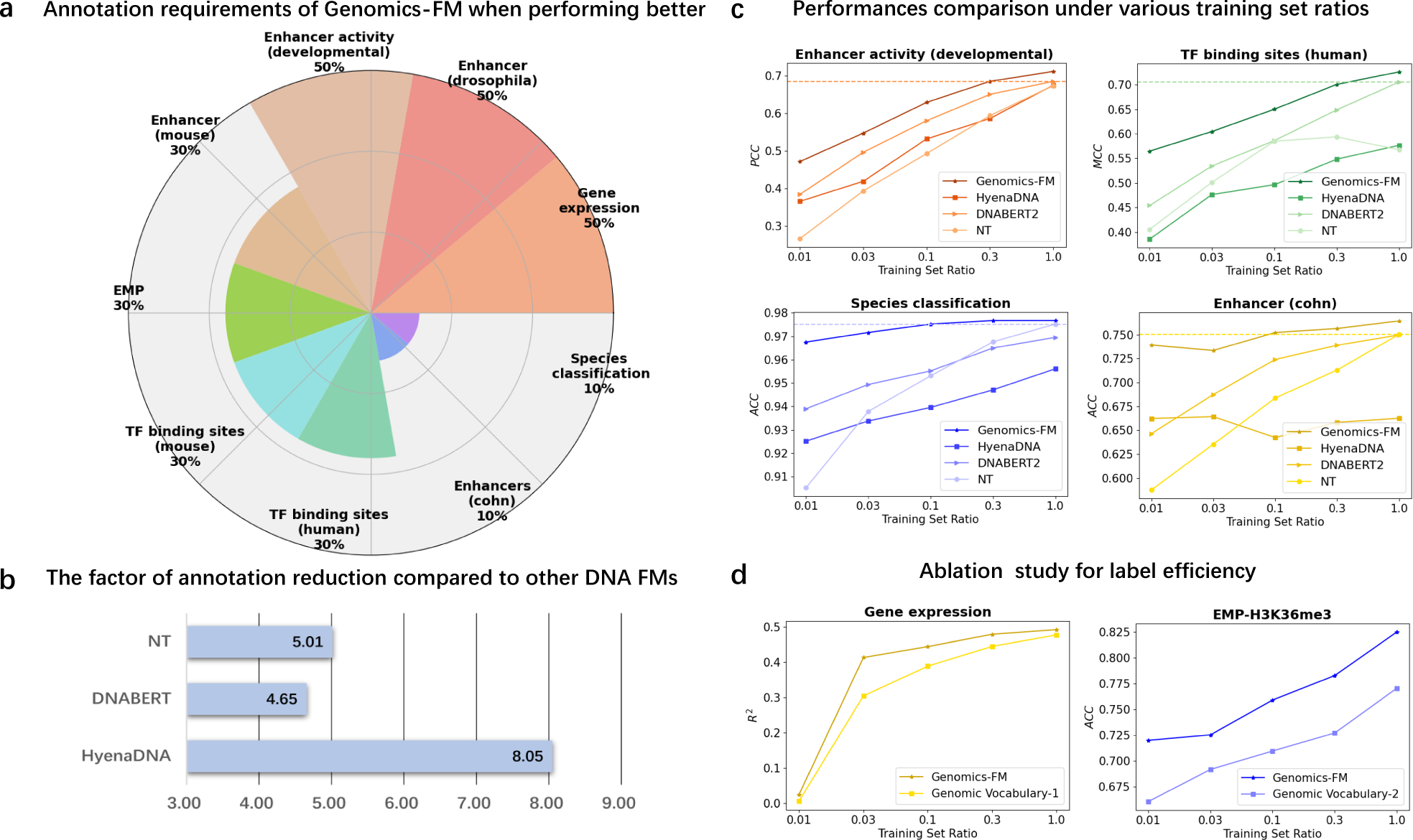
Validation of superior label efficiency of Genomics-FM. **a,** Annotation requirements of Genomics-FM to achieve previous best performances across various tasks, Genomics-FM can achieve the current SOTA performance with much fewer labeled data on a wide range of genetic tasks, greatly reducing the cost of wet-lab experiments. **b,** Mean annotation reduction compared to other DNA foundation models, Genomics-FM reduces the annotations by 5 to 8 times. **c,** Comparisons between Genomics-FM and other DNA foundation models when finetuning under various training set ratios. The dashed line represents the previous best performance finetuned with a full training set. Genomics-FM consistently outperforms other DNA foundation models. **d,** Ablation study for verifying that our vocabulary design can improve label efficiency, Genomics-FM obtains better label efficiency than alternatives that are pretrained and finetuned on a single vocabulary.

#### Comparisons between Genomics-FM and other methods under various training set ratios

We investigate the performance of various DNA foundation models across varying training set ratios, specifically at 1%, 3%, 10%, 30%, 50%, and 100%. The performances are presented in Fig. 3c. Genomics-FM consistently obtains higher performance than other DNA foundation models, underscoring its data efficiency. More impressively, Genomics-FM demonstrates more stable performance when annotations are severely restricted. For example, in the TF binding sites (human) task, with a training set ratio of 1%, Genomics-FM attains an MCC of 0.56 and considerably outperforms the second place (DNABERT2) by 0.11. These results strengthen the potential of Genomics-FM in data-limited applications and scenarios.

#### Influence of the vocabulary design on Label Efficiency

Additional experiments are carried out to determine how the vocabulary design of Genomics-FM contributes to enhancing label efficiency. As depicted in Fig. 3d, in contrast to those models pretrained and finetuned on a single vocabulary, such as Genomic Vocabulary-1 and Genomic Vocabulary-2, our Genomics-FM attains better performance under diverse training set ratios. These results verify that the vocabulary design of Genomics-FM significantly reduces the reliance on labeled samples and substantially improves label efficiency. The main reasons are two-fold: By pretraining with an ensemble vocabulary, Genomics-FM exploits compensatory information from various vocabularies, learning more potent representations; By finetuning with a specific vocabulary, Genomics-FM tailors suitable information for specific tasks, guiding the model to quickly learn task-relevant knowledge.

### 2.4 Genomics-FM enables cross-species prediction

Despite the tremendous achievements in genome sequencing of various species, the epigenomic features such as chromatin accessibility, TF binding, and histone markers remain unclear, apart from a few model organisms like humans and mice. Previous studies have shown that AI models trained on sequences and epigenetic markers of the model organism can be transferred to other species for predictions (Zhang et al., 2022a; Lotfollahi et al., 2022). However, their cross-species prediction performance is still limited due to cross-species data bias (Fithian et al., 2015). To show the superior cross-species predictive capability of Genomics-FM, we construct two datasets: dataset-1 includes histone modification (H3K4me3, H3K27Ac) data from the liver tissues of various mammals, and dataset-2 comprises data from multiple tissues and various epigenetic markers (H3K4me3, H3K27Ac) in three species: pig, cow, chicken. We compare the model under two settings: 1) within-species prediction, where finetuning and testing are performed on data from the same species, and 2) cross-species prediction, where finetuning is conducted on human data and testing is done on data from other species.

#### Cross-species capabilities across various (primate and non-primate) species

As shown in Fig. 4a, Genomics-FM demonstrates exceptional cross-species predictive performance on both primate and non-primate species, surpassing traditional convolutional neural networks (CNNs) in both cross-species and within-species prediction accuracy. We also test the cross-species capabilities of other DNA foundation models, including DNABERT2 and NT (Fig. 4a). For the histone modification H3K4me3, Genomics-FM achieves a cross-species AUC of 0.960, outperforming DNABERT2’s 0.947 and NT’s 0.946. Similarly, for H3K27ac, Genomics-FM records a cross-species AUC of 0.849, exceeding DNABERT2’s 0.819 and NT’s 0.827. Notably, among foundation models, only Genomics-FM’s cross-species prediction surpasses the within-species results of CNNs, which are 0.949 for H3K4me3 and 0.832 for H3K27ac. This enhanced capability is attributed to the strategy of ensemble vocabularies based pretraining, a crucial factor in transcending the typical limitations associated with phylogenetic distance. For non-primate species, although slightly inferior to species-specific models, Genomics-FM demonstrates significant transferability regardless of the phylogenetic distance from humans. We further assess the model’s transfer capabilities by calculating the performance ratio of each model across species relative to its within-species performance. As illustrated in Fig. 4b, while similar performance is observed on primate species closely related to humans, Genomics-FM shows surprisingly better transferability on non-primate species compared to CNN, DNABERT2 and NT.

**Figure 4.**
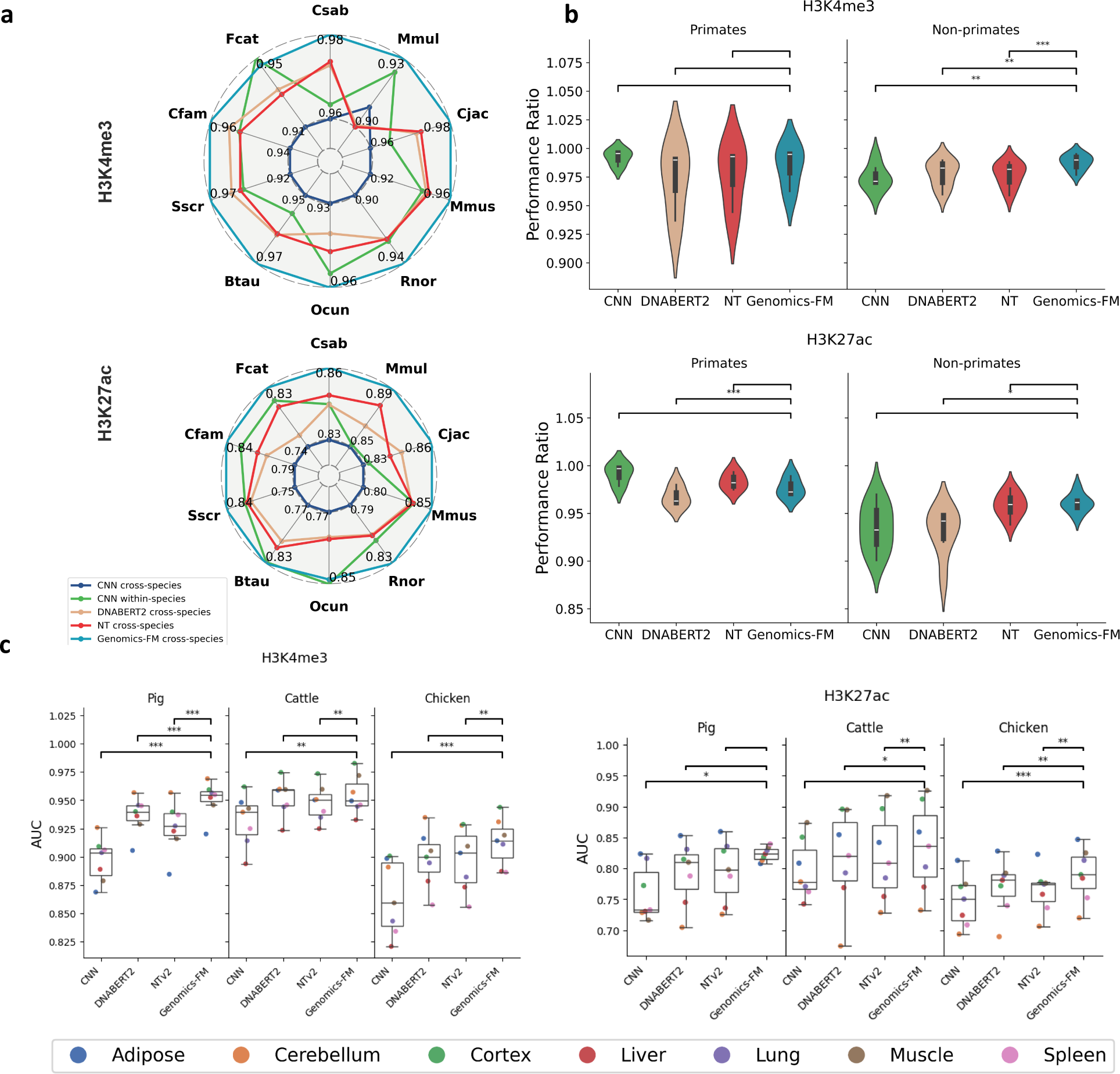
Genomics-FM enables cross-species prediction. **a,** Comparative performance evaluation of different models across ten species with two histone modifications. Genomics-FM shows superior performance compared to CNN in both within-species and cross-species settings. Additionally, Genomics-FM outperforms DNABERT2 and NT in cross-species evaluations. **b,** Evaluation of model transferability across primate and non-primate species. Performance comparisons between Genomics-FM, CNN, DNABERT2, and NT reveal that Genomics-FM exhibits greater improvement over the other models in non-primate species, highlighting its enhanced transferability in species with greater phylogenetic distance from humans. **c,** Cross-species comparison in seven tissues for chicken, pig, and cattle with two histone modification types. Genomics-FM consistently achieves better performance compared to other models.We perform a comparison between Genomics-FM and other model using a two-sided t-test (\**p <* 0.05, \*\**p <* 0.01, \*\*\**p <* 0.001).

#### Cross-species capabilities across various tissues

In dataset-2, we extend our analysis of cross-species capabilities for H3K27Ac and H3K4me3, focusing on multi-tissue comparisons. Each modification type includes data from seven tissues per species. As depicted in Fig.4c, Genomics-FM shows consistent performance improvements compared to other methods across all settings. Due to chickens’ greater phylogenetic distance from humans, their performance is lower than that of pigs and cattle for both modifications. For H3K4me3, CNN’s results vary widely among the seven tissues, with an average AUC of 0.863. Genomics-FM, in contrast, achieves a significantly better and consistent AUC of 0.913 across tissues, indicating robust transfer learning. Genomics-FM also outperforms DNABERT2, which had an average AUC of 0.898. In addition, in the more challenging H3K27Ac analysis, especially with the phylogenetically distant chicken species, Genomics-FM still demonstrates superior and consistent performance across all seven tissues.

### 2.5 Genomics-FM enables cross-tissue prediction

The substantial cost of multi-omics data sequencing and the lack of tissue-specific genomic data pose significant challenges in genomic research. Cross-tissue prediction offers a solution by leveraging existing data to predict omics data for various tissues. To evaluate Genomics-FM’s capability in cross-tissue prediction, we employ the gene expression prediction task and apply our model to predict gene expression levels across different tissues. Although the same DNA sequence is present in all tissues and cell types, specific genes manifest varying expression levels in different cells. Previous studies have indicated that incorporating TF expression data with sequences can enhance the precision of gene expression predictions (Liu et al., 2022). With this understanding, we built the cross-tissue prediction model that incorporates not only DNA sequences but also tissue-specific TF expression data. We collect 1,651 TF expression data from the GTEx database, spanning a diverse array of tissues, after excluding TFs not expressed in any tissue, we retain the tissue-specific expressions of 1,575 TFs. To minimize the impact of external factors such as sequencing depth, we normalize the TF expression data. The gene expression data, sourced from Xpresso (Agarwal & Shendure, 2020), comprise 53 tissues from a consistent origin. These tissues are grouped into 11 categories based on function and region, and the average expression levels are computed to serve as the gene expression label for each tissue group.

#### Effectiveness of incorporating TF expression features

For this task, the sequence data used across all tissues generally remains consistent. Due to normalization effects and intrinsic tissue similarities, even relying solely on sequence data can provide satisfactory predictions of tissue-specific gene expression. However, sequences alone inevitably lack tissue-specific information. By further incorporating transcription factor (TF) expression data, we extract tissue-specific information for each tissue, allowing Genomics-FM to make accurate tissue-specific predictions. As shown in Fig. 5a, the incorporation of TF expression data significantly enhances Genomics-FM’s predictive performance.

**Figure 5.**
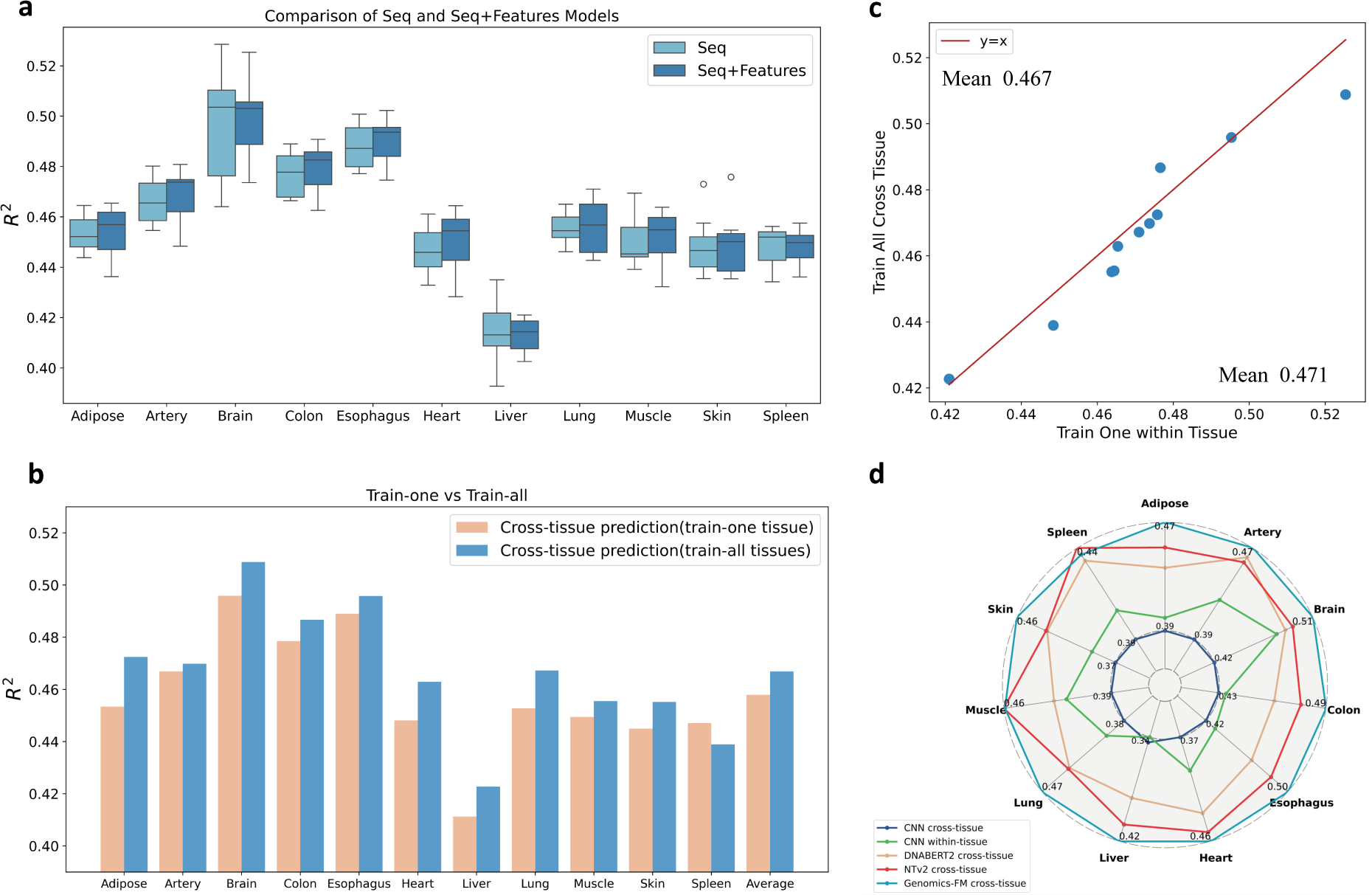
Cross-tissue prediction of Genomics-FM. **a,** The incorporation of transcription factor (TF) expression features can significantly enhance the cross-tissue prediction capability of Genomics-FM beyond what is achievable with sequence data alone. **b,** An examination of the impact of using different tissue data during training, training on multiple tissues (Train-all) improves cross-tissue prediction of Genomics-FM compared to training on a single tissue (Train-one). **c,** Training with data from multiple tissues enhances Genomics-FM’s cross-tissue predictive performance, closely approaching the performance of models directly trained on the target tissue. **d,** A comparison of various methods, Genomics-FM outperforms other methods, achieving the best results.

#### Impact of the integration of multi-tissues

To comprehensively investigate Genomics-FM’s cross-tissue predictive capability, we establish two training settings. “Train-one” involves training the model using data from a single tissue and then predicting in other tissues. “Train-all” entails training the model with data from all tissues except the one being predicted, followed by testing on that specific tissue. As shown in Fig. 5b, the “train-all” setting achieves significantly better cross-tissue prediction performance, the average *R*^2^ value for “train-all” cross-tissue prediction reaches 0.467, higher than the 0.458 achieved by “train-one.” These results indicate that employing a broader array of tissue data rather than focusing on a single tissue significantly boosts the model’s ability to predict across tissues, without increasing computational consumption. Further, as demonstrated in Fig. 5c, by efficiently utilizing data from a broader range of tissues, Genomics-FM’s cross-tissue prediction performance even becomes comparable to the performance achieved when trained directly on the target tissue (*R*^2^ = 0.471).

#### Performance comparison of different methods

Following the “train-all” protocol, we compare the cross-tissue prediction capabilities between Genomics-FM and various models. As illustrated in Fig. 5d, the CNN model’s cross-tissue performance was poor, with an average within-tissue *R*^2^ prediction of only 0.411. In contrast, the NT and DNABERT2 models demonstrate better cross-tissue prediction effectiveness, achieving *R*^2^ values of 0.442 and 0.453, respectively. These foundation models significantly outperform the traditional CNN model in this task. Under the same cross-tissue setting, the Genomics-FM model can achieve the best results, with an average *R*^2^ of 0.467.

### 2.6 Genomics-FM enables cross-molecule prediction

Based on the transcription rules of the central dogma, we conjecture that Genomics-FM, pretrained on large-scale genome sequences, has implicitly acquired knowledge about RNA function and engineering. Thus, we conduct several RNA-specific tasks to evaluate the cross-molecule capabilities of Genomics-FM. For the Mean Ribosome Loading (MRL) prediction task (Sample et al., 2019), which measures translational activity and protein synthesis levels under specific conditions, we follow the protocols used by RNA-FM (Chen et al., 2022) and include datasets from both random (Random Set) and human (Human Set) samples for evaluation. As engineered RNA elements function as programmable detectors for small molecules, proteins, and nucleic acids, predicting their behavior is both important and challenging in synthetic biology, thus we utilize the BEACON benchmark (Ren et al., 2024) to predict the state activity of RNA switches. To investigate alternative polyadenylation (APA), a prevalent regulatory mechanism in gene expression across various organisms, we predict the proximal APA isoform ratio (Isoform) for each variant in the 3’UTR sequence using different libraries from APARENT (Bogard et al., 2019).

#### Comparison between Genomics-FM and RNA models under RNA tasks

As shown in Fig. 6a, we compare Genomics-FM with traditional SOTA RNA methods and the powerful RNA language model RNA-FM. First, Genomics-FM can outperform traditional SOTA RNA methods across different tasks. For example, for the 5’UTR function, Genomics-FM achieves an *R*^2^ score of 0.825, surpassing the SOTA method Optimus-5-prime (Sample et al., 2019) which scores 0.78. For the 3’UTR function, Genomics-FM attains the *R*^2^ score of 0.7319, exceeding the SOTA method APARENT (Bogard et al., 2019) which scores 0.5082. Second, Genomics-FM also outperforms the powerful open-source RNA foundation model RNA-FM. For instance, Genomics-FM obtains an *R*^2^ score of 0.5760 in RNA switches prediction, surpassing RNA-FM which scores 0.5598. It is worth noting that RNA-FM is pretrained on approximately 23 million non-coding RNA data, whereas Genomics-FM is pretrained on DNA data and is not explicitly trained on RNA data in the pretraining phase. This indicates that, despite being trained on long genomic sequences without explicit sequence annotations, Genomics-FM still demonstrates an understanding of constitutive RNA sequences.

**Figure 6.**
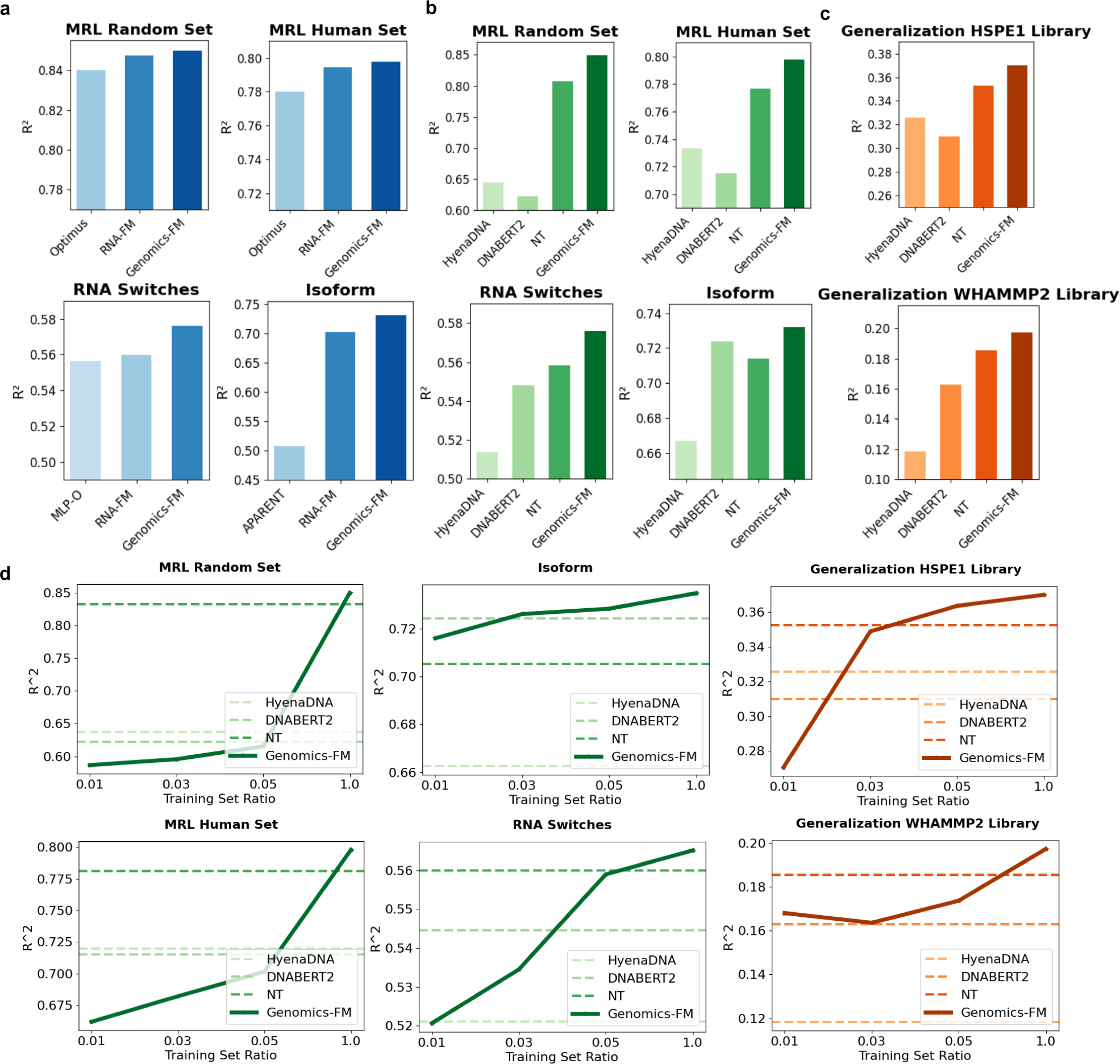
Cross-molecule capacities of Genomics-FM for RNA tasks. a, Cross-molecule ability validation of Genomics-FM on three important RNA tasks including Mean Ribosome Loading, RNA switches and APA Isoform prediction, Genomics-FM outperforms both traditional RNA methods and popular RNA foundation model. b, Comparison between Genomics-FM and other DNA foundation models on RNA tasks, Genomics-FM consistently outperforms its peers in various complex RNA prediction scenarios. c, Generalization ability validation of different DNA foundation models on unseen RNA libraries, Genomics-FM shows robust performance across diverse datasets, confirming its adaptability and effectiveness in handling unseen RNA sequences. d, Label efficiency of DNA foundation models on RNA tasks, Genomics-FM achieves significant performance gains with fewer labeled data.

#### Comparison between Genomics-FM and DNA foundation models under RNA tasks

As shown in Fig. 6b, our Genomics-FM demonstrates superior cross-molecule capability compared to other advanced DNA foundation models, including HyenaDNA, DNABERT2, and Nucleotide Transformer (NT). Specifically, on the MRL random set, Genomics-FM achieves a score of 0.8498, significantly outperforming HyenaDNA, which scored 0.6441, DNABERT2 with 0.6223, and Nucleotide Transformer at 0.8075. These results suggest that our model excels at learning the implicit knowledge of the central dogma during training and mining the transcriptional functions of the genome during pretraining, thereby enhancing the understanding of RNA functions.

#### Generalization of Genomics-FM across unseen RNA libraries

Moreover, in order to analyze the ability of Genomics-FM to efficiently understand RNA functionality, we collect different RNA libraries from the MPRAs database to evaluate its zero-shot capability. We train the model on several databases and then test it on unseen databases HSPE1 and WHAMMP2 (Bogard et al., 2019). Fig. 6c shows that Genomics-FM exhibits superior generalization performance. In the HSPE1 Library, Genomics-FM achieves an *R*^2^ value of 0.37, outperforming other models such as HyenaDNA and DNABERT2, which reach only 0.3258 and 0.3098, respectively. Similarly, in the WHAMMP2 Library, Genomics-FM leads with an *R*^2^ value of 0.1974, significantly ahead of the closest competitor, Nucleotide Transformer, which scores about 0.1856. This demonstrates that Genomics-FM has substantial potential to predict and discover new RNA functions.

#### Label efficiency analysis of Genomics-FM on RNA tasks

We compare the performance of various methods under different training set ratios. As shown in Fig. 6d, Genomics-FM makes more efficient use of RNA data compared to other DNA foundation models, even if RNA sequences were not explicitly seen during pretraining. Specifically, when using 100% of the data, the deep green and brown curves representing Genomics-FM consistently outperform other methods such as HyenaDNA, DNABERT2, and Nucleotide Transformer. When using a few RNA samples, Genomics-FM has the potential to achieve comparability with other models with a mass of data. For example, Genomics-FM finetuned with only 1% data can outperform HyenaDNA with 100% data on the unseen WHAMMP2 library. These results suggest that Genomics-FM presents novel opportunities for advancing research in mRNA-based therapy and RNA design.

## 3 Discussion

Decoding the genomic language is a longstanding and unresolved field, characterized by an abundance of unlabeled data, diverse tasks, and a scarcity of labeled samples. In this paper, we introduce Genomics-FM, a versatile and label-efficient functional genomic foundation model. Recognizing the unique characteristics of different genomic data and tasks, Genomics-FM employs an ensemble vocabulary during pre-training to extract richer feature representations from vast amounts of unlabeled data, and utilizes a specific vocabulary during fine-tuning to better capture task-specific features from limited labeled data. Genomics-FM achieves new state-of-the-art performance across 16 tasks spanning regulatory element annotation, gene expression and chromatin state, variations, and species-level analyses, and significantly reduces the annotation requirements needed to achieve the performance of existing state-of-the-art models, which is critically important for resource-constrained biological research. Impressively, Genomics-FM has proven exceptionally effective in various cross-domain scenarios, including cross-species, cross-tissue, and cross-molecule analyses. For cross-species prediction tasks, Genomics-FM outperforms both conventional models and other DNA foundation models, even achieving predictive capabilities comparable to within-species predictions of conventional models, with its transferability remaining robust even with increasing evolutionary distance. Similarly, in cross-tissue prediction tasks, Genomics-FM’s performance is on par with within-tissue predictions. Additionally, Genomics-FM excels in cross-molecule analyses, demonstrating strong generalization to RNA prediction tasks and achieving performance comparable even superior to widely-used RNA foundation models.

Despite these advancements in various tasks, Genomics-FM still faces some limitations in the current version. Although benefits from an ensemble vocabulary are illustrated, Genomics-FM utilizes only two vocabularies during the pretraining stage. From the technical perspective, it is straightforward to expand our framework to incorporate more mainstream vocabularies, which will be presented in our subsequent version. Additionally, rather than using vocabulary construction techniques from the natural language processing field, more endeavors will be devoted to developing biologically meaningful vocabularies. Besides, although we have provided insights into the way in which vocabulary is selected for downstream tasks, Genomics-FM still lacks vocabulary automatic selection capabilities during the finetuning stage. One possible solution is to leverage embeddings from multiple vocabularies concurrently during the fine-tuning phase, the advantage is that it maximizes diverse encoding information from various vocabularies, while the drawback is that it does not consider the specific characteristics of the tasks. Future work could delve deeper into developing task-agnostic vocabulary selection strategies and providing more biological interpretability. Additionally, the power of scaling law in Genomics-FM remains underexplored. Experiment results have shown the benefits when scaling up the pretrained data volume and model size, and we will subsequently explore the impact of larger models and larger data on the representation learning of DNA sequences.

Looking ahead, Genomics-FM plans to expand its capabilities in several key areas to explore new frontiers in genomic research. One focus will be on enhancing multi-tasking abilities, especially for diverse DNA sequence-based tasks such as gene expression, structural analysis, and functional assessments of noncoding regions. Another critical area is improving label efficiency in more scenarios where annotated data are scarcer, which could transform the model’s utility in under-resourced biological settings. We also aim to extend our model’s applicability across species by incorporating more diverse prediction categories and developing a cross-species mapping to deepen our understanding of genomic similarities and differences across various organisms. Furthermore, efforts will be made to broaden the scope of tasks beyond gene expression, encompassing a wider range of biological tissues and molecular tasks. Finally, we are looking to advance our model’s capabilities in handling RNA-related tasks and eventually tackle protein sequence-based challenges, which hold significant promise for creating a unified representation of biological sequences across different modalities. These planned enhancements are crucial for enabling Genomics-FM to support a broader range of genomic research objectives, thereby facilitating the development of more comprehensive and versatile tools in the field of genomics.

## A Supplemental material

### A.1 Model

#### A.1.1 Model Architecture

In our research, we have employed the BERT structure from the transformer model as our foundational pre-trained model, which adopts an encoder-only architecture. This model utilizes Masked Language Modeling (MLM) for supervision. The input length of the model is constrained to a maximum of 512 tokens. To enhance length extrapolation in downstream tasks, we have incorporated Attention with Linear Biases (ALiBi), a form of relative position encoding. This approach enables our model to handle input sequence lengths in downstream tasks that exceed the length of the pre-training data.

#### A.1.2 Tokenizer and Data Preparation

To cater to varying lengths and granularities in downstream tasks, we trained two types of tokenizers during the pre-training phase: BPE (Byte Pair Encoding) and KMER.

BPE is widely used in natural language processing and operates on a frequency-based statistical approach. In the context of DNA sequences, BPE gradually incorporates high-frequency nucleotide combinations into the vocabulary until it reaches a predetermined size. BPE is a non-overlapping tokenizer, segmenting the input sequence according to the vocabulary. Longer nucleotide combinations present in the vocabulary are prioritized as tokens. Given its ability to encode multiple nucleotides per token and its non-overlapping segmentation, BPE can encode longer nucleotide sequences for the same input length, albeit not at the granularity of individual nucleotides. This indicates that BPE can achieve longer sequence lengths compared to KMER.

KMER functions as a fixed-length tokenizer, encoding ‘K’ nucleotides per token. We employ an overlapping KMER approach, where the window size is ‘K’, and the step is 1. This tokenizer can achieve single nucleotide granularity, as consecutive tokens overlap by ‘K-1’ nucleotides. However, KMER cannot handle excessively long downstream tasks. For our research, we selected the commonly used 6mer variant.

#### A.1.3 Data Preparation

Different tokenizer methods necessitate distinct approaches to input data processing due to their varying capacities to encode nucleotides. Moreover, to adapt to different lengths in downstream tasks, we dynamically segment DNA sequences to obtain gene sequences of varying lengths.

For BPE, we control the length of input DNA sequences within a range of 70-2300, with sequences of 2300 nucleotides constituting 50% and the remaining range accounting for the other 50%. For 6mer, the length is controlled between 70-510, with 510 nucleotide sequences comprising 50% and other lengths making up the remaining 50%.

For input DNA sequences, we use BPE and 6mer as tokenizers, converting them into tokens for input data. We add CLS, SEP, and some special prompt tokens as inputs to the model. The inputs are subjected to varying degrees of masking. For inputs generated by the BPE tokenizer, excluding special tokens, a subset of tokens is randomly selected for masking. Of these, 80% are replaced with MASK tokens, 10% remain unchanged, and another 10% are randomly replaced with other tokens. For 6mer, due to its overlapping segmentation, at least six consecutive tokens need to be masked to prevent information leakage. The model is then tasked with predicting the masked tokens. Through this methodology, the model learns the contextual relationships in DNA sequences, thereby enhancing predictions in downstream tasks.

### A.2 Pretrain Dataset

In order to accommodate a broader range of downstream tasks, our pre-training data encompassed multi-species data as well as data from the 1000 Genomes Project (1KG).

For the multi-species data, we referenced approaches from Nucleotide and GENA-LM, primarily sourcing from NCBI and Ensembl. We selected at least one species from each category, culminating in a comprehensive dataset spanning 1,100 species across diverse classifications including Fungi, Bacteria, Protozoa (Invertebrates), Mammalian, Birds, Reptiles, Amphibians, and Fish. To prevent loss of genetic information due to sequence trimming, we implemented an overlapping scheme (100 nucleotide overlap), thereby accumulating a total of 284 billion nucleotides. Thanks to the broad coverage of our pre-trained multi-species dataset, our model demonstrated exceptional performance in cross-species tasks.

Regarding the 1KG data, we followed the processing method of GENA-LM, with data derived from gnomAD. This dataset comprised variant genomes from 38 individuals around the globe, representing 19 distinct ethnicities, with an equal gender distribution. For each genomic region of these gnomAD samples, sequences were generated through allelic replacements, utilizing information from all SNP loci, resulting in 385 billion nucleotides. The inclusion of variant data not only empowered our model to excel in tasks involving genetic variations but also enhanced performance in human-related tasks. This is attributed to the 1KG data serving as a significant data augmentation for human genetic information.

### A.3 Benchmark Dataset

#### A.3.1 Epigenetic Marks Prediction

The dataset we used, which was previously compiled and identified in the yeast genome, focuses on epigenetic marks, specifically acetylation and methylation nucleosome occupancies. These occupancies were initially determined through Chip-Chip experiments and were subsequently converted into datasets categorizing them as either positive or negative occurrences. This conversion was essential for creating a training dataset for epigenetic analysis. The dataset includes a range of histone marks such as H3, H4, H3K9ac, H3K14ac, H4ac, H3K4me1, H3K4me2, H3K4me3, H3K36me3, and H3K79me3.

#### A.3.2 Promoter 300 Detection

The study of promoter detection in humans primarily involves identifying the (proximal) promoter regions in the human genome. These regions are essential for initiating transcription and contain key regulatory elements. Understanding these sites is crucial for gaining insights into gene regulation and identifying genetic factors related to various diseases. The dataset used in this research is categorized into two types: TATA and non-TATA promoters, depending on the presence of a TATA box motif within the sequence. For each category, we extract sequences spanning from -249 to +50 base pairs around the Transcription Start Site (TSS). These sequences are sourced from the Eukaryotic Promoter Database (EPDnew). They form our ‘promoter class’. Conversely, the ‘non-promoter class’ is composed of sequences of equal length randomly chosen from areas outside the promoter regions. These include sequences with a TATA motif (termed TATA non-promoters) and sequences without a TATA box (non-TATA, non-promoters). Additionally, we merge the TATA and non-TATA promoter datasets to create a comprehensive dataset, referred to as ‘all’.

#### A.3.3 Promoter Core Detection

Human core promoter detection shares similarities with proximal promoter detection but specifically targets predicting the core promoter region. This region, central and closest to the Transcription Start Site (TSS) and the start codon, is crucial for transcription initiation. Unlike proximal promoter detection, core promoter detection involves a much narrower context window, typically ranging from -34 to +35 base pairs around the TSS. This narrower focus makes core promoter detection a more complex and challenging task compared to proximal promoter prediction.

#### A.3.4 Transcription Factor Binding Site Prediction (Human)

The task of predicting human transcription factor binding sites involves identifying where transcription factors (TFs), essential proteins that regulate gene expression, bind in the human genome. Accurately predicting these sites is crucial for understanding genetic interactions and identifying targets for gene therapy. For this study, we utilized data from 690 legacy ENCODE ChIP-seq experiments, as found in the UCSC genome browser. This data includes 161 TF binding profiles across 91 human cell lines. We focused on a 101-base pair region centered around each peak, classifying this as the TFBS (Transcription Factor Binding Site) class. For the non-TFBS class, we selected non-overlapping sequences of the same length and GC content. To refine our analysis, we chose 5 datasets from a subset of the original 690. This selection was based on heuristic filtering, where we excluded tasks that were either too simple (with an F1 score above 0.95) or too difficult (with an F1 score below 0.50) for current language models.

#### A.3.5 Splice Site Prediction

The focus of human splice site prediction is to identify the precise locations of splice donor and acceptor sites in the human genome. These sites are key to alternative splicing, a process central to protein diversity and understanding the role of aberrant splicing in genetic disorders. Our dataset, derived from the Ensembl GRCh38 human reference genome, features sequences 400 base pairs in length. Previous models, as indicated by (Ji et al., 2021). in 2021, have shown near-perfect performance on the original dataset, which contains 10,000 sequences each of splice donors, acceptors, and non-splice sites. However, this level of accuracy is overly optimistic for detecting non-canonical sites in real-world applications. To address this, we enhanced the dataset by adding adversarial examples. These are sequences that represent unseen false positive predictions from a hold-out set, thus introducing a higher level of complexity and challenge to the task.

#### A.3.6 Transcription factor binding site prediction (Mouse)

The goal of mouse transcription factor binding site prediction is to identify where transcription factors attach to the mouse genome. This process is akin to the human binding site prediction. We utilized mouse ENCODE ChIP-seq data from the UCSC genome browser, which, as of the latest data by Stamatoyannopoulos et al. in 2012, includes 78 datasets. Unlike the human TFBS prediction, where negative examples were generated differently, in the mouse version, negative examples are created through dinucleotide shuffling. This method maintains the relative frequencies of the base pairs. All other aspects of the dataset preparation mirror the approach used for the human TFBS prediction. Additionally, we applied the same selection method as in the human study, randomly choosing 5 datasets out of the total 78 available, based on our predetermined criteria.

#### A.3.7 Chromatin Profiles Prediction

In our research on chromatin profiles prediction, we utilized a dataset originally compiled by (Zhou & Troyanskaya, 2015). This dataset consists of 2.4 million sequences, each 1000 nucleotides long, linked to 919 different chromatin features. These features encompass 690 transcription factors (TFs), 125 DNAse features, and 104 histone characteristics. Our model was designed to handle these 919 classification tasks simultaneously, equipped with 919 separate classification heads. We calculated the loss as the mean of the cross-entropy losses across these tasks. Given the imbalance in each label, predominantly composed of negative samples, we amplified the losses for positive samples by a factor of eight. This approach differs from the DeepSEA method, which trained two separate models—one for forward sequences and another for reverse-complementary sequences—and then averaged their predictions. In contrast, our model was trained exclusively on the forward sequences.

#### A.3.8 Drosophila Enhancers Prediction

We acquired candidate sequences and their corresponding data on housekeeping and tissue-specific activities in Drosophila cells from the Stark Lab repository. These datasets were divided into training, validation, and testing sets, following the same partitioning used in training the DeepSTARR model. Our research involves a two-class regression task. In this task, each sequence, 249 base pairs in length, is analyzed to predict two continuous scores: the first score indicates the level of housekeeping enhancer activity, and the second score reflects the developmental enhancer activity.

#### A.3.9 Enhancer(cohn)

The Cohn dataset for Human enhancers, adapted from (Cohn et al., 2018), presents enhancers as genomic regulatory functional elements. These elements have the capability to bind with specific DNA binding proteins, thereby regulating the transcription of targeted genes. Distinct from promoters, enhancers are not required to be in close proximity to the genes they affect. In fact, they can be located several million bases away. This significant distance poses a challenge in accurately detecting enhancers.

#### A.3.10 Enhancer Design

##### Developmental Enhancers

These enhancers are crucial for regulating gene expression during the development of an organism. They ensure genes are expressed at the right time and place during development.

##### Housekeeping Enhancers

These enhancers regulate genes essential for maintaining basic cellular functions, typically expressed across all cell types and developmental stages.

The datasets for studying developmental and housekeeping enhancer activities typically involve high-throughput sequencing techniques such as UMI-STARR-seq. This method allows for the generation of quantitative activity maps of enhancers across the genome. For example, in the DeepSTARR project (de Almeida et al., 2022), UMI-STARR-seq was used to create high-resolution maps of enhancer activity in Drosophila melanogaster S2 cells. This dataset includes genome-wide mea-surements of both developmental and housekeeping enhancer activities, providing a comprehensive resource for enhancer research.

#### A.3.11 Coding region

The coding vs non-coding task within the Genomic Benchmark dataset aims to classify genomic sequences as either protein-coding or non-coding. This task is crucial for understanding the functional elements of the genome and differentiating between sequences that produce proteins and those that perform regulatory or other non-coding roles.The datasets used for this task include sequences that have been carefully curated and preprocessed to ensure high quality and relevance. The positive class consists of sequences known to be protein-coding, while the negative class includes non-coding sequences. Each sequence is labeled and divided into training and testing subsets to facilitate model development and evaluation.

#### A.3.12 the Human Enhancers Ensemb

In the case of the Human enhancers Ensembl dataset, construction was based on enhancers from The FANTOM5 project (Andersson et al., 2014), accessed via the Ensembl database (Howe et al., 2021). This dataset features negative sequences randomly generated from the Human genome GRCh38. These sequences match the lengths of positive sequences and are designed to not overlap with them.

#### A.3.13 The Human Regulatory Ensembl

The Human regulatory Ensembl dataset, also derived from the Ensembl database (Howe et al., 2021), encompasses three classes: enhancer, promoter, and open chromatin region, as identified in The Ensembl Regulatory Build (Hoskins et al., 2007). The open chromatin region sequences in this dataset align with the positive sequences found in the Human ocr Ensembl dataset.

#### A.3.14 Spieces classification

The species classification task primarily distinguishes between human and worm (C. elegans) genomic sequences. The human vs worm task in the Genomic Benchmark dataset is primarily designed to distinguish between genomic sequences from humans and worms (C. elegans). This classification task utilizes the unique genomic signatures inherent to each species, enhancing our understanding of species-specific genomic features and their functional implications.The dataset comprises genomic sequences sourced from both humans and C. elegans. These sequences are labeled accordingly and split into training and testing subsets to support the development and evaluation of machine learning models. This task facilitates comparative genomic studies by enabling the precise classification of sequences from different species, thereby aiding in the creation of species-specific genomic tools and the exploration of evolutionary biology.

#### A.3.15 Variant Prediction

We categorized the evaluation of variant effects into two settings: fine-tune and zero-shot prediction.

##### Fine-Tune Setting

The fine-tune process, illustrated in Fig. S1, involves several steps. Initially, the model is trained on chromatin profiling tasks, resulting in a chromatin spectrum model. This model is then used to extract features from the data before and after the introduction of variants. The differences between these features serve as inputs for the second stage, where supervised fine-tuning is conducted. For this process, the dataset is divided into training and test sets to evaluate the model’s performance.

##### Zero-Shot Prediction Setting

The zero-shot prediction process, shown in Fig. S2, also starts with fine-tuning the model using chromatin profiling data. The fine-tuned model is then employed to extract features from the sequences of the entire dataset, both before and after variant introduction. These features are used to compute the distances between the pre- and post-variant sequences using four different calculation methods. Finally, Pearson correlation is utilized to assess the variant effects, providing a measure of the impact of the variants without additional fine-tuning.

### A.4 Label efficiency

To validate the label efficiency of our model, we processed the downstream task data from the aforementioned benchmarks. Specifically, we randomized the data and sequentially saved 1%, 3%, 10%, 30%, 40%, 50%, 60%, 70%, 80%, and 90% of the dataset to simulate scenarios with scarce labeled data. This method ensures that larger subsets of the dataset include all smaller subsets, thereby reducing variability due to data fluctuations to some extent. Additionally, the validation and test datasets remain unchanged, consistent with the full data finetuning process of the task. The hyperparameters for finetuning on the new datasets are also maintained as consistent as possible to verify the model’s label efficiency capability.

### A.5 Cross-Species

#### Data collection

Genome versions corresponding to the datasets were downloaded from the UCSC Genome Browser (https://hgdownload.soe.ucsc.edu). We introduce three settings and corresponding genome versions as follows.

1. We selected histone modification data for 11 mammals (human, rhesus monkey, vervet monkey, marmoset, mouse, rat, rabbit, cow, pig, dog and cat) (Villar et al., 2015) as shown in Table S6. The dataset includes two types of modifications: H3K4me3 and H3K27ac. The data is sourced from liver tissues and comes from E-MTAB-2633. The downloaded data does not require preprocessing.
2. We chose multi-omics sequencing data from three species (chicken, pig, and cattle) and seven different tissues (adipose, cerebellum, cortex, liver, lung, muscle, spleen) (Kern et al., 2021; Pan et al., 2021) as shown in Table S7. The data is from GSE158430. Additionally, we specifically selected CTCF, H3K27ac, H3K4me3, and chromatin accessibility for further analysis. For each species and tissue, we calculated the intersection of repeated peaks using the dba.peakset function from DiffBind (Stark et al., 2011), with the minOverlap parameter set to 0.999. This process is applied to all biological replicates corresponding to each species and tissue.
3. Due to the absence of epigenetic data for human tissues in the references corresponding to Setting II, we downloaded sequencing data for human tissues and epigenetic types from the ENCODE database (Sloan et al., 2016) (https://www.encodeproject.org/) as shown in Table S8. The downloaded data does not require preprocessing.

#### Division of Positive and Negative Sets

We split the data samples into positive and negative sets.

1. Positive Set: Complete Chromatin Open Regions (OCR) corresponding to peaks.
2. Negative Set: Random sampling from the intergenic regions of OCR. The number of peaks in the negative set is controlled to be equal to the positive set in a 1:1 ratio. Additionally, the negative set is controlled to have consistent chromosome, sequence length, and G-C content (G-C content difference less than 1%) with the positive set.

### A.6 Cross-tissue

**Gene expression** is fundamental to understanding the functional dynamics within transcriptionally active regions of the genome. It provides essential insights into how genetic information is converted into cellular structures and functions. Specifically, gene expression analysis reveals variations in mRNA and protein levels across different tissues and conditions, illustrating the complex interplay of genetic and environmental factors.

Our study utilizes a comprehensive collection of gene and TF expression datasets. We extracted 1651 expression datasets from the GTEx database, covering a wide range of tissues. After filtering out TFs not expressed in any sampled tissue, we retained 1575 TF expression datasets. These datasets underwent normalization to reduce the influence of external factors like sequencing depth. For gene expression data, we used data from Xpresso (Agarwal & Shendure, 2020), which included consistent measurements across the first 53 tissues. These tissues were classified into 11 functional and regional groups, and average expression levels were calculated to label each tissue group. This meticulous data organization supports our model’s ability to produce reliable and accurate predictions. The evaluation of the gene expression task employs the *R*^2^ value as the key metric, which quantifies the proportion of variance in cross-tissue gene expression that our model can accurately predict.

### A.7 Cross-molecule

**Mean Ribosome Loading (MRL)** serves as a predictive measure for the translational activity of mRNA sequences into proteins, quantified by MRL values. Our analysis computes MRL across a robust dataset of 91,519 sequences from the 5’ untranslated region (UTR) and their variants, derived from Reid’s extensive collection (Sample et al., 2019). The precision of our predictive model is validated through the coefficient of determination, or *R*^2^ value, which assesses the predictability of variance in the dependent variable based on the independent variables.

**Programmable RNA Switches (PRS)** focuses on the synthesis and functional adaptation of RNA molecules, which can dynamically alter their structure in response to specific external signals. The operational states of these RNA sequences—ON, OFF, and ON/OFF—are quantitatively assessed using the metric *y* ∈ R^3^. The dataset under analysis by (Angenent-Mari et al., 2020) encompasses 91,534 toehold switch sequences in vivo, spanning 23 viral genomes and 906 human transcription factors. The activity levels of these switches are indicated through GFP signal intensities for both ON and OFF states (Angenent-Mari et al., 2020). The efficacy of PRS is rigorously evaluated using the *R*^2^ metric, highlighting the predictive power of our model.

**Alternative Polyadenylation Isoform Prediction (APA)** employs predictive analytics to determine the usage ratios of proximal polyadenylation sites (PAS) within the 3’ untranslated region (3’ UTR) of each genetic variant, with results captured in the target variable *y* ∈ R. This analysis filters and examines 228,000 sequences from a substantial repository of over three million APA (Bogard et al., 2019) reporter gene data entries found in Bogard’s dataset. The task is structured as a regression model, focusing on the proportion of proximal APA isoforms. Model performance is quantified using the *R*^2^ metric, which measures the accuracy of the predictions.

### A.8 Model Analysis

#### Vocabulary analysis

Selecting an appropriate vocabulary for downstream tasks can lead to performance improvements. In our study, we employed multiple vocabularies during pre-training to accommodate diverse requirements across downstream tasks. Given the distinct preferences for vocabulary among different tasks, we selectively fine-tuned specific vocabularies based on task characteristics. We proposed a straightforward yet effective selection criterion: we derived a task-specific vocabulary by extracting statistical information from input sequences (positive samples for binary classification tasks), then intersected it with the pre-training vocabulary to identify the vocabulary with the highest overlap for fine-tuning.

Experimental results demonstrate that among the 14 downstream tasks investigated, 11 tasks conform to this criterion. We conducted testing on all tasks using both KMER and BPE vocabularies, revealing distinct preferences for tokenizer methods across tasks. As depicted in Fig. S3, among the 7 tasks where KMER exhibited superior performance, 6 tasks showed a higher overlap between their task-specific vocabularies and the KMER vocabulary. Similarly, among the 7 tasks where BPE performed better, 5 tasks exhibited a higher overlap between their task-specific vocabularies and the BPE vocabulary. Thus, when multiple pre-trained vocabularies are available, and testing on all of them is not desired, selecting vocabularies with higher overlap is beneficial, as their performance tends to be superior in most cases.

#### Scaling decipherment

This part sheds light on the performance improvements on DNA downstream tasks through scaling of the model, pre-train data, and vocabularies. Scaling decipherment observed in large language models have proven effective in DNA pretraining as well. For instance, in Nucleotide Transformer models, scaling up both the model and data has shown promising results. We corroborate this observation through experiments, and further validating the efficacy of prompt training in our model. As depicted in Fig. S3b, leveraging multi-species data resulted in performance enhancements across the majority of tasks compared to models trained solely on the human reference genome, with an average improvement of 0.6%. Concurrently, scaling up the model size also yielded performance gains in downstream tasks. Large models with 400 million parameters, as opposed to base models with 120 million parameters, exhibited an average improvement of 0.4%. Our prompt training model, with the same parameter count (400 million), achieves an additional performance improvement of 0.6%. All of these findings underscore the significance of scaling up both the model and data. Furthermore, they highlight that selecting an appropriate vocabulary can further enhance performance in practical applications.

#### Biological interpretation

Our model has successfully identified unique sequence patterns in DNA. In biology, a motif refers to short sequence fragments with distinct functions, commonly found in proteins or DNA/RNA sequences, such as protein-binding sites on DNA. Accurately recognizing these motifs is crucial for assessing the cognitive abilities of the model in the field of biology. Using a previously developed motif analysis tool DNABERT-viz (Ji et al., 2021), we conducted an in-depth analysis of the attention weights of the finely tuned model. By applying the hypergeometric distribution hypothesis test, we successfully identified several specific DNA motifs, including BNC2, Mlxip, fos-1, PIF1, and JDP2. The model’s predictive results, as shown in Fig5c, are highly consistent with the actual laboratory sequencing results in Jaspar (Rauluseviciute et al., 2024). These findings demonstrate that the model, trained on extensive data, does not merely memorize but effectively focuses on the sequence patterns in DNA through its attention mechanism and delves deeply into their biological features.

## B Supplementary Figures

**Figure S1.**
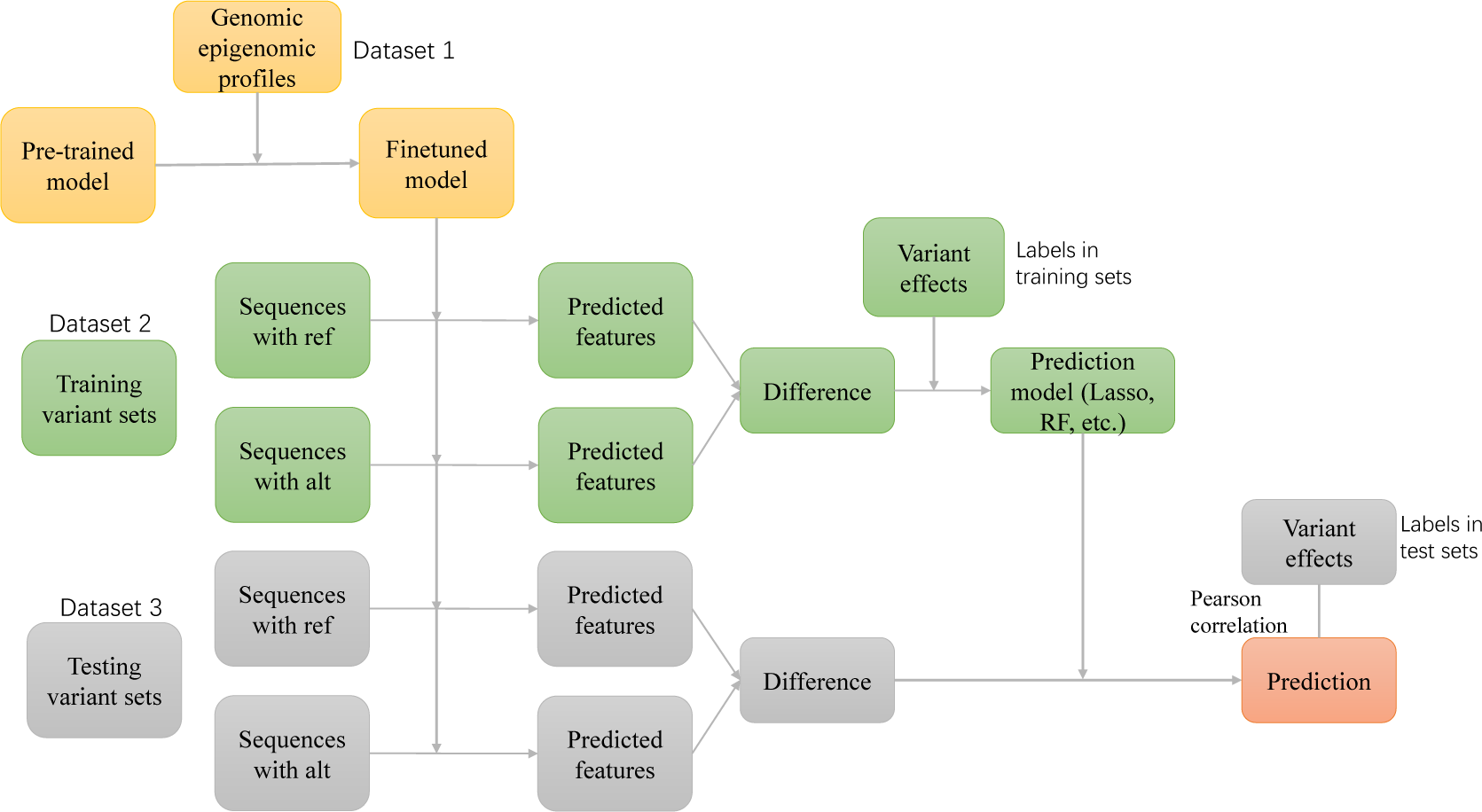
Pipeline of supervised variant effect prediction.

**Figure S2.**
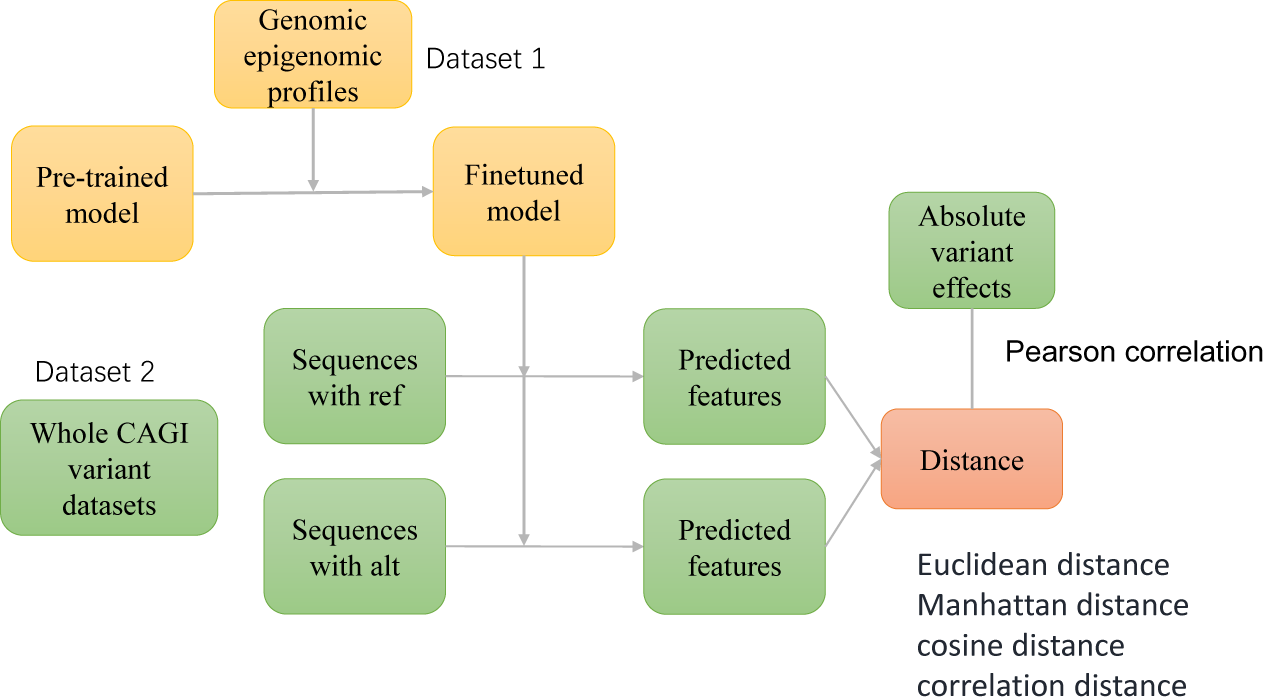
Pipeline of zero-shot variant effect prediction.

**Figure S3.**
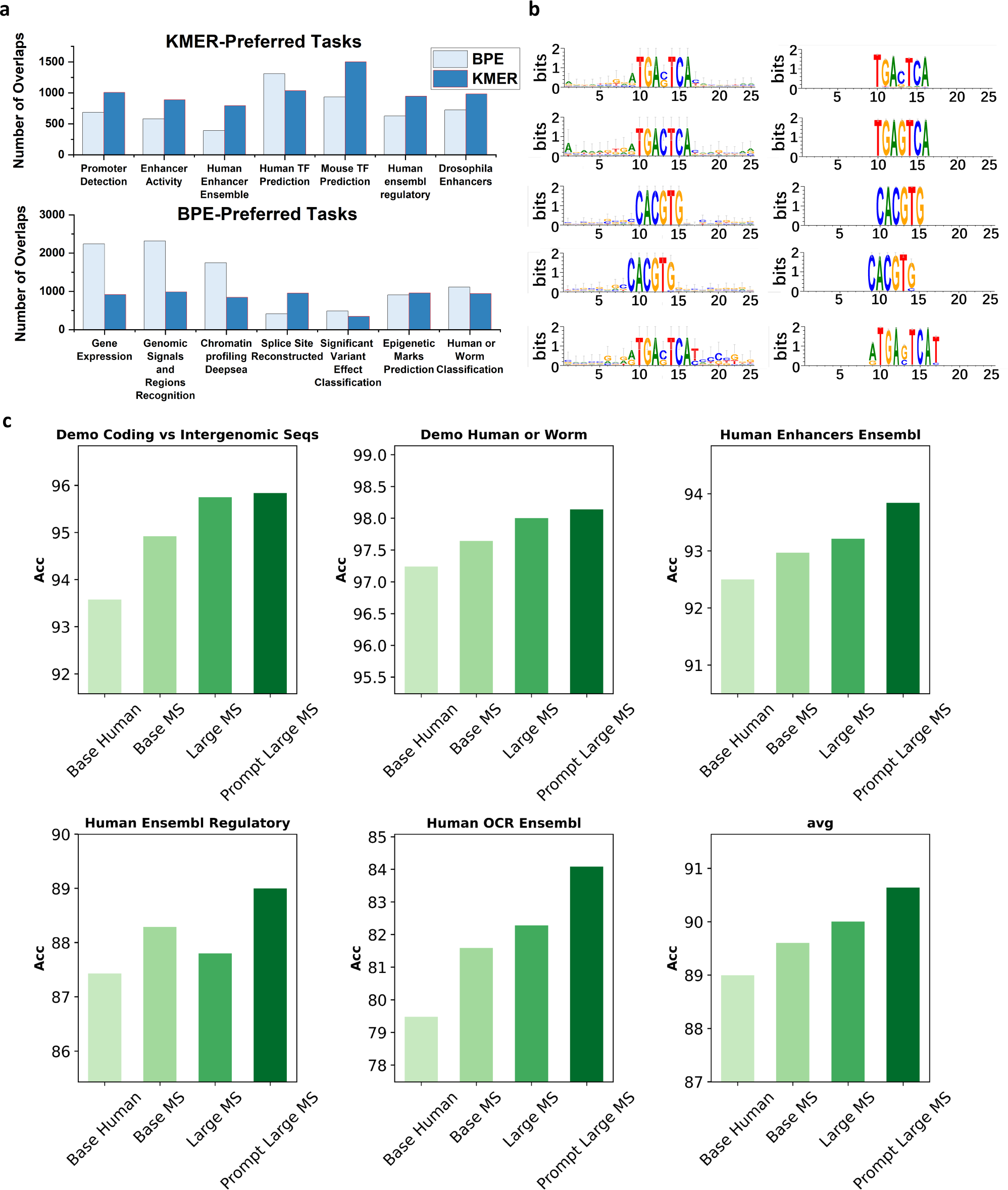
Model Analysis. a. Overlap between the vocabulary of BPE and KMER token sets with task-specific BPE tokens derived from positive samples in downstream tasks. The top chart highlights tasks where KMER performed better, while the bottom chart shows tasks where BPE excelled. b. Comparison of predicted motifs from the model with those in the JASPAR 2024 database. c. Scaling up the model and data consistently improved performance across most tasks. Additionally, our prompt model further enhanced performance.

## C Supplementary Tables

**Table S1.**
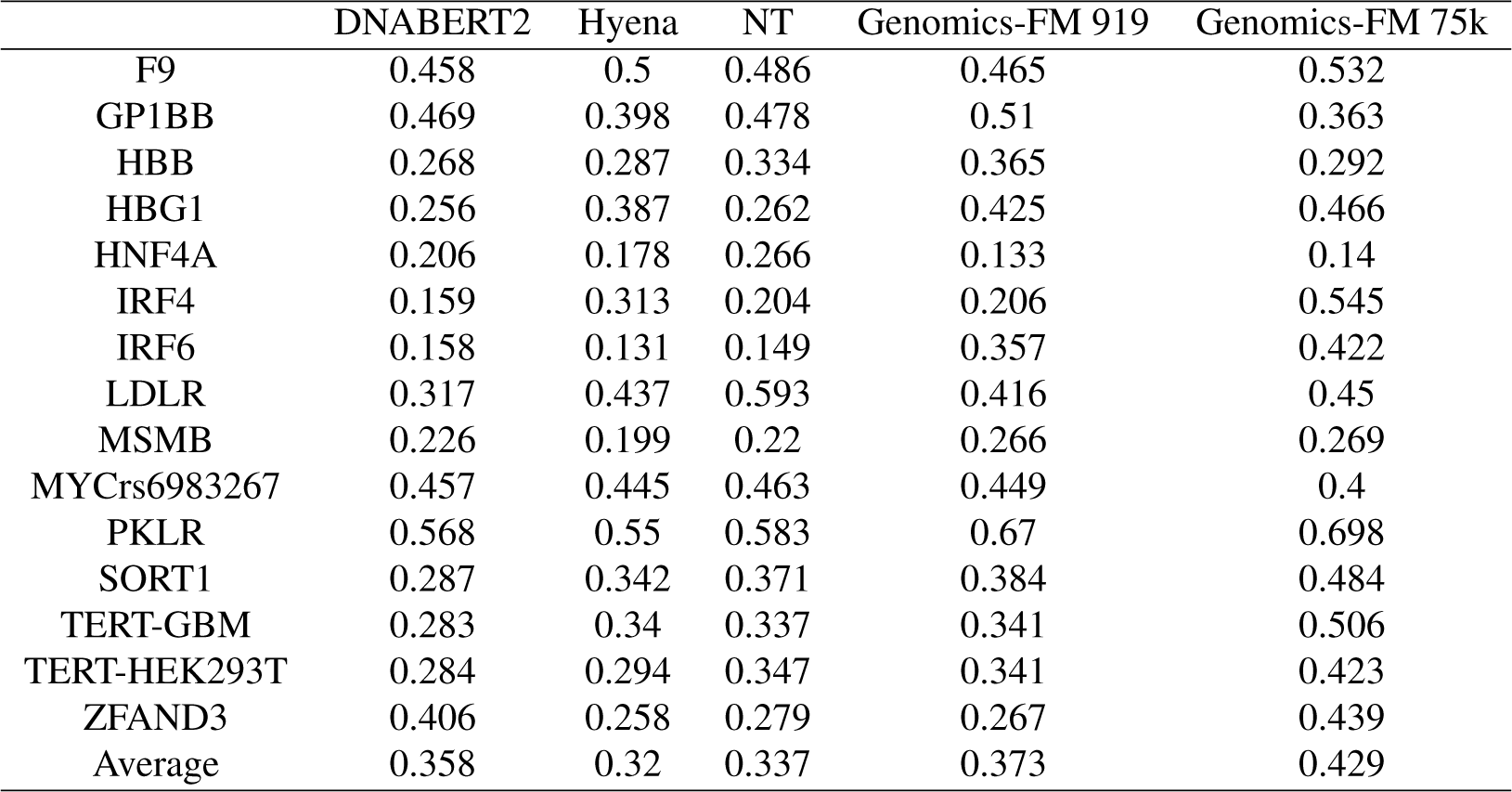
Euclidean-Based Evaluation of Variant Effects in finetuning Prediction Models.

|  | DNABERT2 | Hyena | NT | Genomics-FM 919 | Genomics-FM 75k |
| --- | --- | --- | --- | --- | --- |
| F9 | 0.458 | 0.5 | 0.486 | 0.465 | 0.532 |
| GP1BB | 0.469 | 0.398 | 0.478 | 0.51 | 0.363 |
| HBB | 0.268 | 0.287 | 0.334 | 0.365 | 0.292 |
| HBG1 | 0.256 | 0.387 | 0.262 | 0.425 | 0.466 |
| HNF4A | 0.206 | 0.178 | 0.266 | 0.133 | 0.14 |
| IRF4 | 0.159 | 0.313 | 0.204 | 0.206 | 0.545 |
| IRF6 | 0.158 | 0.131 | 0.149 | 0.357 | 0.422 |
| LDLR | 0.317 | 0.437 | 0.593 | 0.416 | 0.45 |
| MSMB | 0.226 | 0.199 | 0.22 | 0.266 | 0.269 |
| MYCr6983267 | 0.457 | 0.445 | 0.463 | 0.449 | 0.4 |
| PKLR | 0.568 | 0.55 | 0.583 | 0.67 | 0.698 |
| SORT1 | 0.287 | 0.342 | 0.371 | 0.384 | 0.484 |
| TERT-GBM | 0.283 | 0.34 | 0.337 | 0.341 | 0.506 |
| TERT-HEK293T | 0.284 | 0.294 | 0.347 | 0.341 | 0.423 |
| ZFAND3 | 0.406 | 0.258 | 0.279 | 0.267 | 0.439 |
| Average | 0.358 | 0.32 | 0.337 | 0.373 | 0.429 |

**Table S2.** Euclidean-Based Evaluation of Variant Effects in Zero-Shot Prediction Models.

|  | DNABERT2 | Hyena | NT | Genomics-FM 919 | Genomics-FM 75k |
| --- | --- | --- | --- | --- | --- |
| F9 | 0.533 | 0.258 | 0.466 | 0.555 | 0.653 |
| GP1BB | 0.357 | 0.311 | 0.403 | 0.564 | 0.397 |
| HBB | 0.521 | 0.42 | 0.447 | 0.537 | 0.403 |
| HBG1 | 0.583 | 0.627 | 0.569 | 0.658 | 0.619 |
| HNF4A | 0.028 | 0.132 | 0.223 | 0.265 | 0.317 |
| IRF4 | 0.407 | 0.448 | 0.207 | 0.393 | 0.797 |
| IRF6 | 0.372 | 0.114 | 0.352 | 0.687 | 0.640 |
| LDLR | 0.304 | 0.382 | 0.578 | 0.513 | 0.337 |
| MSMB | 0.334 | 0.209 | 0.305 | 0.389 | 0.362 |
| MYCr6983267 | 0.511 | 0.311 | 0.49 | 0.469 | 0.509 |
| PKLR | 0.756 | 0.658 | 0.689 | 0.810 | 0.752 |
| SORT1 | 0.337 | 0.472 | 0.456 | 0.527 | 0.558 |
| TERT-GBM | 0.572 | 0.541 | 0.57 | 0.698 | 0.661 |
| TERT-HEK293T | 0.414 | 0.47 | 0.563 | 0.599 | 0.587 |
| ZFAND3 | 0.457 | 0.42 | 0.567 | 0.402 | 0.625 |
| Average | 0.432 | 0.385 | 0.459 | 0.538 | 0.548 |

**Table S3.** Manhattan-Based Evaluation of Variant Effects in Zero-Shot Prediction Models.

|  | DNABERT2 | Hyena | NT | Genomics-FM 919 | Genomics-FM 7w5 |
| --- | --- | --- | --- | --- | --- |
| F9 | 0.49 | 0.524 | 0.52 | 0.477 | 0.529 |
| GP1BB | 0.469 | 0.405 | 0.497 | 0.55 | 0.341 |
| HBB | 0.292 | 0.324 | 0.368 | 0.387 | 0.288 |
| HBG1 | 0.26 | 0.444 | 0.289 | 0.447 | 0.46 |
| HNF4A | 0.231 | 0.213 | 0.293 | 0.145 | 0.137 |
| IRF4 | 0.167 | 0.313 | 0.218 | 0.216 | 0.555 |
| IRF6 | 0.164 | 0.141 | 0.152 | 0.402 | 0.462 |
| LDLR | 0.335 | 0.467 | 0.602 | 0.454 | 0.442 |
| MSMB | 0.258 | 0.218 | 0.244 | 0.31 | 0.27 |
| MYCr6983267 | 0.463 | 0.453 | 0.473 | 0.466 | 0.399 |
| PKLR | 0.59 | 0.58 | 0.607 | 0.707 | 0.719 |
| SORT1 | 0.302 | 0.364 | 0.397 | 0.418 | 0.512 |
| TERT-GBM | 0.306 | 0.399 | 0.356 | 0.356 | 0.499 |
| TERT-HEK293T | 0.3 | 0.333 | 0.365 | 0.349 | 0.418 |
| ZFAND3 | 0.405 | 0.294 | 0.302 | 0.293 | 0.459 |
| Average | 0.379 | 0.335 | 0.365 | 0.398 | 0.433 |

**Table S4.** Cosine-Based Evaluation of Variant Effects in Zero-Shot Prediction Models.

|  | DNABERT2 | Hyena | NT | Genomics-FM 919 | Genomics-FM 75k |
| --- | --- | --- | --- | --- | --- |
| F9 | 0.481 | 0.484 | 0.5 | 0.461 | 0.522 |
| GP1BB | 0.376 | 0.359 | 0.409 | 0.429 | 0.303 |
| HBB | 0.115 | 0.091 | 0.16 | 0.247 | 0.176 |
| HBG1 | 0.072 | 0.188 | 0.081 | 0.251 | 0.388 |
| HNF4A | 0.162 | 0.121 | 0.252 | 0.045 | 0.188 |
| IRF4 | 0.131 | 0.253 | 0.117 | 0.112 | 0.378 |
| IRF6 | 0.117 | 0.103 | 0.126 | 0.249 | 0.269 |
| LDLR | 0.262 | 0.399 | 0.564 | 0.413 | 0.426 |
| MSMB | 0.086 | 0.076 | 0.094 | 0.143 | 0.117 |
| MYCr6983267 | 0.471 | 0.463 | 0.443 | 0.422 | 0.451 |
| PKLR | 0.507 | 0.456 | 0.508 | 0.636 | 0.561 |
| SORT1 | 0.302 | 0.35 | 0.378 | 0.335 | 0.442 |
| TERT-GBM | 0.157 | 0.222 | 0.207 | 0.248 | 0.432 |
| TERT-HEK293T | 0.178 | 0.212 | 0.219 | 0.258 | 0.36 |
| ZFAND3 | 0.283 | 0.202 | 0.155 | 0.112 | 0.32 |
| Average | 0.281 | 0.247 | 0.265 | 0.291 | 0.356 |

**Table S5.** Dot Product-Based Evaluation of Variant Effects in Zero-Shot Prediction Models.

|  | DNABERT2 | Hyena | NT | Genomics-FM 919 | Genomics-FM 75k |
| --- | --- | --- | --- | --- | --- |
| F9 | 0.439 | 0.492 | 0.489 | 0.458 | 0.505 |
| GP1BB | 0.442 | 0.371 | 0.386 | 0.461 | 0.353 |
| HBB | 0.145 | 0.127 | 0.191 | 0.259 | 0.248 |
| HBG1 | 0.189 | 0.222 | 0.148 | 0.296 | 0.535 |
| HNF4A | 0.138 | 0.123 | 0.173 | 0.028 | 0.124 |
| IRF4 | 0.106 | 0.228 | 0.097 | 0.119 | 0.475 |
| IRF6 | 0.132 | 0.075 | 0.099 | 0.271 | 0.248 |
| LDLR | 0.261 | 0.268 | 0.538 | 0.42 | 0.542 |
| MSMB | 0.132 | 0.109 | 0.134 | 0.179 | 0.191 |
| MYCrS6983267 | 0.454 | 0.416 | 0.424 | 0.42 | 0.446 |
| PKLR | 0.498 | 0.45 | 0.525 | 0.638 | 0.58 |
| SORT1 | 0.208 | 0.288 | 0.253 | 0.323 | 0.405 |
| TERT-GBM | 0.2 | 0.121 | 0.209 | 0.255 | 0.482 |
| TERT-HEK293T | 0.205 | 0.123 | 0.22 | 0.263 | 0.377 |
| ZFAND3 | 0.258 | 0.146 | 0.16 | 0.107 | 0.34 |
| Average | 0.27 | 0.254 | 0.237 | 0.3 | 0.39 |

**Table S6.**
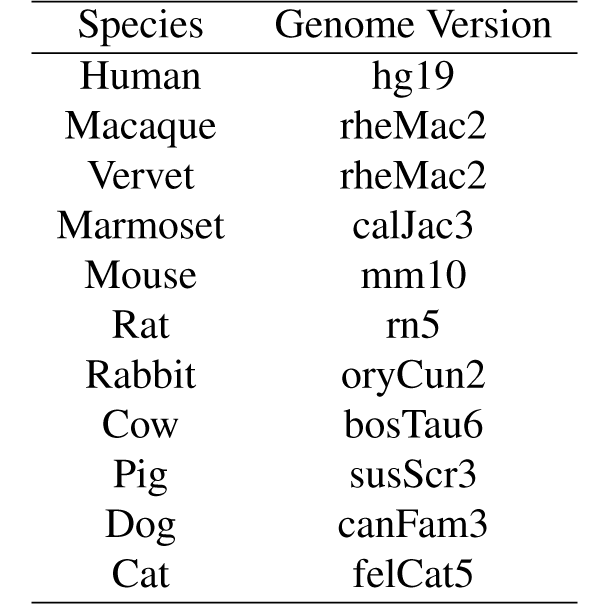
Genome versions corresponding to species in Setting I.

| Species | Genome Version |
| --- | --- |
| Human | hg19 |
| Macaque | rheMac2 |
| Vervet | rheMac2 |
| Marmoset | calJac3 |
| Mouse | mm10 |
| Rat | rn5 |
| Rabbit | oryCun2 |
| Cow | bosTau6 |
| Pig | susScr3 |
| Dog | canFam3 |
| Cat | felCat5 |

**Table S7.** Genome versions corresponding to species in Setting II.

| Species | Genome Version |
| --- | --- |
| Pig | susScr11 |
| Cattle | bosTau9 |
| Chicken | galGal6 |

**Table S8.** Genome version for human in ENCODE.

| Species | Genome Version |
| --- | --- |
| Human | hg38 |

**Table S9.**
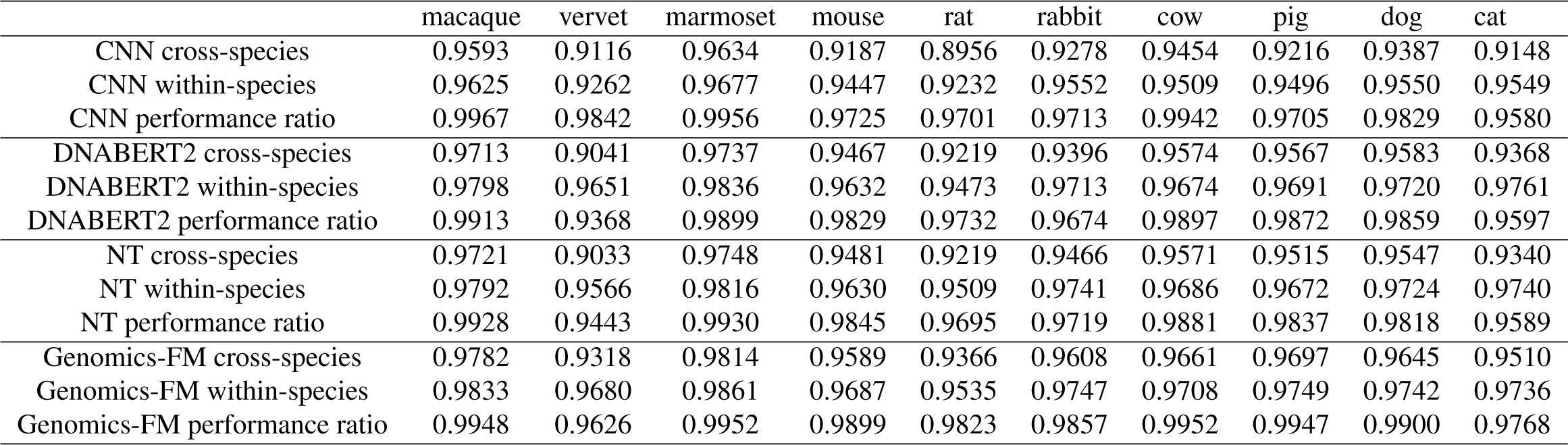
Comparison of model performance on H3K4me3 modification (Dataset 1).

**Table S10.**
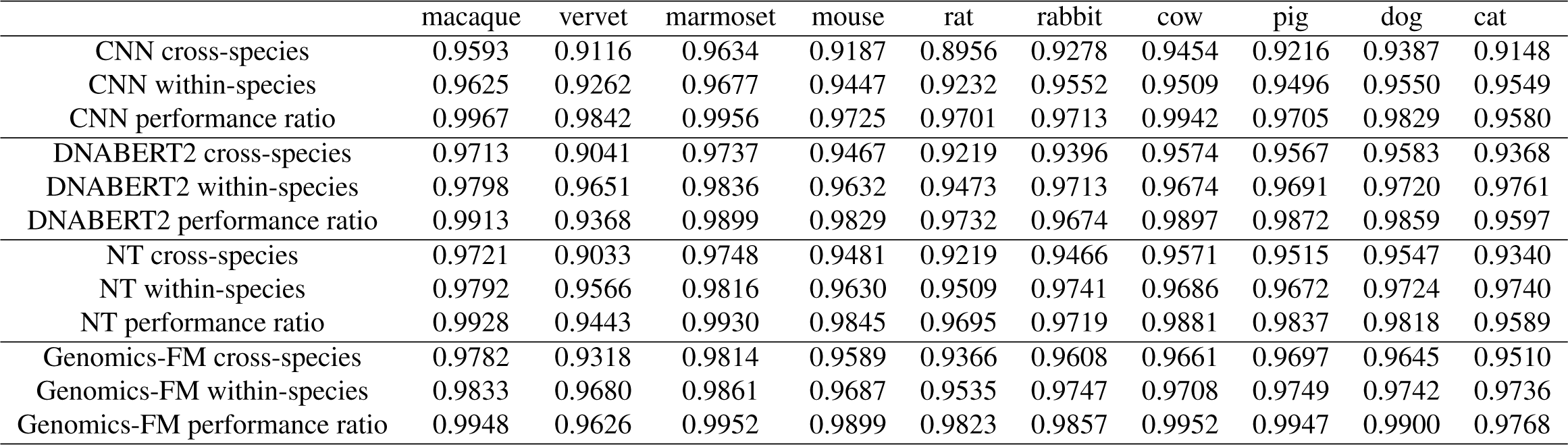
Comparison of model performance on H3K4me3 modification (Dataset 1).

**Table S11.**
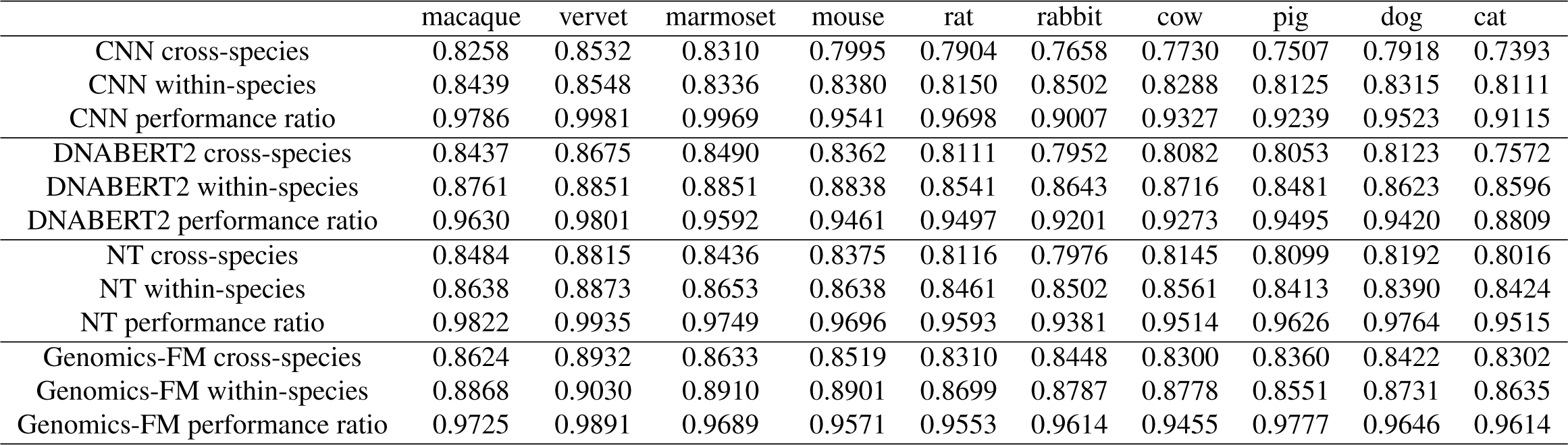
Comparison of model performance on H3K27ac modification (Dataset 1).

**Table S12.**
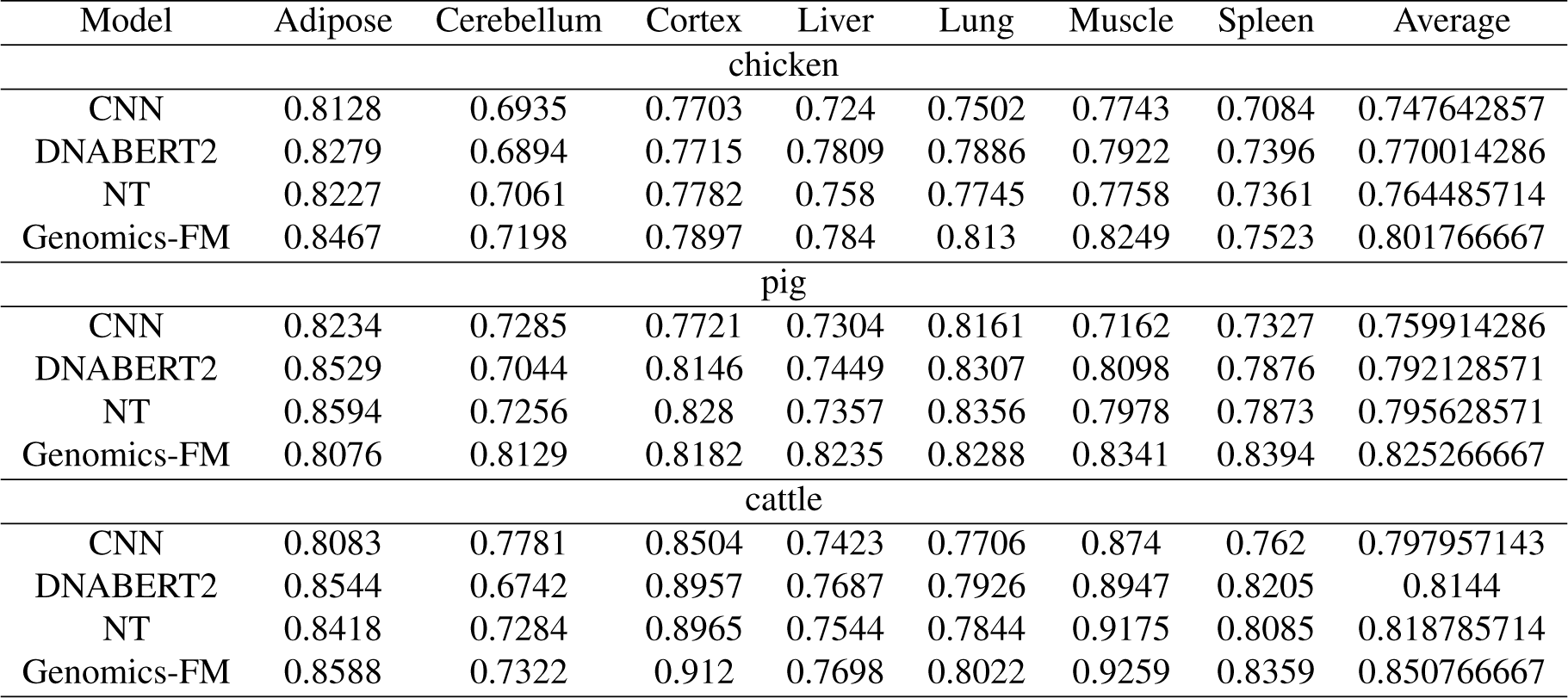
Comparison of model performances on the H3K27ac modification.

**Table S13.** Predictive performance of the genomics-FM model using sequence data for gene expression across various tissues. Diagonal values represent within-tissue prediction accuracy.

| Model | Adipose | Cerebellum | Cortex | Liver | Lung | Muscle | Spleen | Average |
| --- | --- | --- | --- | --- | --- | --- | --- | --- |
| chicken |  |  |  |  |  |  |  |  |
| CNN | 0.8984 | 0.8908 | 0.9007 | 0.8206 | 0.8431 | 0.8592 | 0.8341 | 0.8638 |
| DNABERT2 | 0.9162 | 0.9347 | 0.8998 | 0.8784 | 0.8945 | 0.9052 | 0.8574 | 0.898028571 |
| NT | 0.9033 | 0.9278 | 0.9287 | 0.8731 | 0.882 | 0.9093 | 0.8557 | 0.897128571 |
| Genomics-FM | 0.9141 | 0.9308 | 0.9437 | 0.8871 | 0.911 | 0.919 | 0.8862 | 0.9131 |
| pig |  |  |  |  |  |  |  |  |
| CNN | 0.8687 | 0.9258 | 0.9091 | 0.8889 | 0.9036 | 0.8787 | 0.906 | 0.8973 |
| DNABERT2 | 0.9055 | 0.9567 | 0.9399 | 0.936 | 0.9455 | 0.9287 | 0.9451 | 0.936771429 |
| NT | 0.8845 | 0.9567 | 0.9396 | 0.9226 | 0.9272 | 0.9158 | 0.9371 | 0.926214286 |
| Genomics-FM | 0.9201 | 0.969 | 0.9593 | 0.9524 | 0.9563 | 0.9457 | 0.9548 | 0.9511 |
| cattle |  |  |  |  |  |  |  |  |
| CNN | 0.948 | 0.9394 | 0.9619 | 0.8937 | 0.9141 | 0.9426 | 0.9249 | 0.932085714 |
| DNABERT2 | 0.9588 | 0.9597 | 0.9746 | 0.9233 | 0.9441 | 0.9594 | 0.9458 | 0.952242857 |
| NT | 0.9502 | 0.9506 | 0.9735 | 0.9248 | 0.9347 | 0.9596 | 0.9401 | 0.947642857 |
| Genomics-FM | 0.9494 | 0.9571 | 0.9826 | 0.9326 | 0.9445 | 0.9718 | 0.9458 | 0.9548 |

**Table S14.** Classification of tissue groups by organ and function.

| group | tissue |
| --- | --- |
| Brain | Brain_Amygdala, Brain_Anterior_cingulate_cortex_BA24,<br>Brain_Caudate_basal_ganglia, Brain_Cerebellar_Hemisphere,<br>Brain_Cerebellum, Brain_Cortex, Brain_Frontal_Cortex_BA9,<br>Brain_Hippocampus, Brain_Hypothalamus,<br>Brain_Nucleus_accumbens_basal_ganglia,<br>Brain_Putamen_basal_ganglia, Brain_Spinal_cord_cervical_c1,<br>Brain_Substantia_nigra |
| Heart | Heart_Atrial_Appendage, Heart_Left_Ventricle |
| Artery | Artery_Aorta, Artery_Coronary, Artery_Tibial |
| Liver | Liver |
| Colon | Colon_Sigmoid, Colon_Transverse |
| Esophagus | Esophagus_Gastroesophageal_Junction, Esophagus_Mucosa, Esophagus_Muscularis |
| Lung | Lung |
| Skin | Skin_Not_Sun_Exposed_Suprapubic, Skin_Sun_Exposed_Lower_leg |
| Adipose | Adipose_Subcutaneous, Adipose_Visceral_Omentum |
| Spleen | Spleen |
| Muscle | Muscle_Skeletal |

**Table S15.** Predictive Performance of the Genomics-FM Model Using Sequence Data for Gene Expression Across Various Tissues. Diagonal Values Indicate Within-Tissue Prediction Accuracy.

|  | Adipose | Artery | Brain | Colon | Esophagus | Heart | Liver | Lung | Muscle | Skin | Spleen |
| --- | --- | --- | --- | --- | --- | --- | --- | --- | --- | --- | --- |
| Adipose | 0.4630 | 0.4734 | 0.5046 | 0.4875 | 0.4974 | 0.4533 | 0.4225 | 0.4620 | 0.4556 | 0.4579 | 0.4565 |
| Artery | 0.4643 | 0.4764 | 0.5026 | 0.4886 | 0.4985 | 0.4607 | 0.4268 | 0.4678 | 0.4612 | 0.4592 | 0.4585 |
| Brain | 0.4667 | 0.4819 | 0.5048 | 0.4920 | 0.5025 | 0.4642 | 0.4275 | 0.4662 | 0.4665 | 0.4555 | 0.4556 |
| Colon | 0.4688 | 0.4799 | 0.5013 | 0.4889 | 0.4995 | 0.4641 | 0.4230 | 0.4718 | 0.4643 | 0.4594 | 0.4588 |
| Esophagus | 0.4642 | 0.4790 | 0.5126 | 0.4892 | 0.5009 | 0.4580 | 0.4150 | 0.4643 | 0.4606 | 0.4579 | 0.4537 |
| Heart | 0.4716 | 0.4829 | 0.5175 | 0.4978 | 0.5082 | 0.4639 | 0.4303 | 0.4738 | 0.4634 | 0.4638 | 0.4631 |
| Liver | 0.4788 | 0.4890 | 0.5148 | 0.5012 | 0.5137 | 0.4719 | 0.4356 | 0.4810 | 0.4705 | 0.4732 | 0.4697 |
| Lung | 0.4679 | 0.4808 | 0.5109 | 0.4932 | 0.5034 | 0.4608 | 0.4256 | 0.4676 | 0.4636 | 0.4622 | 0.4630 |
| Muscle | 0.4715 | 0.4815 | 0.5029 | 0.4949 | 0.5062 | 0.4650 | 0.4293 | 0.4786 | 0.4604 | 0.4633 | 0.4629 |
| Skin | 0.4617 | 0.4730 | 0.5116 | 0.4863 | 0.4960 | 0.4531 | 0.4191 | 0.4599 | 0.4544 | 0.4570 | 0.4527 |
| Spleen | 0.4653 | 0.4764 | 0.5081 | 0.4892 | 0.4987 | 0.4580 | 0.4231 | 0.4704 | 0.4578 | 0.4570 | 0.4583 |

**Table S16.** Predictive performance of the genomics-FM model using sequence data incorporating with transcription factor expression features for gene expression across various tissues. Diagonal values represent within-tissue prediction accuracy.

|  | Adipose | Artery | Brain | Colon | Esophagus | Heart | Liver | Lung | Muscle | Skin | Spleen |
| --- | --- | --- | --- | --- | --- | --- | --- | --- | --- | --- | --- |
| Adipose | 0.4724 | 0.4851 | 0.5083 | 0.4965 | 0.5069 | 0.4681 | 0.4320 | 0.4721 | 0.4716 | 0.4680 | 0.4622 |
| Artery | 0.4594 | 0.4698 | 0.4996 | 0.4812 | 0.4954 | 0.4532 | 0.4196 | 0.4619 | 0.4578 | 0.4591 | 0.4595 |
| Brain | 0.4606 | 0.4725 | 0.5088 | 0.4858 | 0.4960 | 0.4515 | 0.4125 | 0.4577 | 0.4513 | 0.4479 | 0.4483 |
| Colon | 0.4619 | 0.4750 | 0.5138 | 0.4867 | 0.4991 | 0.4556 | 0.4161 | 0.4607 | 0.4572 | 0.4549 | 0.4530 |
| Esophagus | 0.4625 | 0.4732 | 0.5038 | 0.4857 | 0.4958 | 0.4582 | 0.4151 | 0.4616 | 0.4563 | 0.4573 | 0.4530 |
| Heart | 0.4663 | 0.4808 | 0.5170 | 0.4920 | 0.5051 | 0.4629 | 0.4197 | 0.4659 | 0.4644 | 0.4569 | 0.4527 |
| Liver | 0.4718 | 0.4857 | 0.5216 | 0.4959 | 0.5100 | 0.4678 | 0.4227 | 0.4703 | 0.4657 | 0.4637 | 0.4566 |
| Lung | 0.4664 | 0.4796 | 0.5038 | 0.4913 | 0.5039 | 0.4609 | 0.4232 | 0.4672 | 0.4646 | 0.4665 | 0.4647 |
| Muscle | 0.4609 | 0.4734 | 0.5045 | 0.4863 | 0.4972 | 0.4546 | 0.4266 | 0.4644 | 0.4555 | 0.4590 | 0.4589 |
| Skin | 0.4630 | 0.4708 | 0.5048 | 0.4874 | 0.4959 | 0.4571 | 0.4170 | 0.4670 | 0.4538 | 0.4552 | 0.4535 |
| Spleen | 0.4535 | 0.4651 | 0.5110 | 0.4806 | 0.4878 | 0.4443 | 0.4066 | 0.4523 | 0.4436 | 0.4460 | 0.4389 |

**Table S17.** Comparison of Genomics-FM model performance under train-one and train-all settings. Train-one involves training the model on individual tissues separately, while train-all uses all tissues collectively for training.

|  | Adipose | Artery | Brain | Colon | Esophagus | Heart | Liver | Lung | Muscle | Skin | Spleen |
| --- | --- | --- | --- | --- | --- | --- | --- | --- | --- | --- | --- |
| Train-one cross-tissue | 0.4534 | 0.4669 | 0.4959 | 0.4785 | 0.4890 | 0.4481 | 0.4112 | 0.4527 | 0.4494 | 0.4449 | 0.4470 |
| Train-one within-tissue | 0.4654 | 0.4738 | 0.5254 | 0.4766 | 0.4953 | 0.4644 | 0.4210 | 0.4710 | 0.4638 | 0.4758 | 0.4484 |
| Train-all cross-tissue | 0.4724 | 0.4698 | 0.5088 | 0.4867 | 0.4958 | 0.4629 | 0.4227 | 0.4672 | 0.4555 | 0.4552 | 0.4389 |

**Table S18.** Comparison of model performance under the train-all setting and seq + features setting. The train-all setting involves training the model on all available tissues collectively, whereas the seq + features setting incorporate sequence data with additional TF expression features for enhanced prediction accuracy.

|  | Adipose | Artery | Brain | Colon | Esophagus | Heart | Liver | Lung | Muscle | Skin | Spleen |
| --- | --- | --- | --- | --- | --- | --- | --- | --- | --- | --- | --- |
| CNN-cross-tissue | 0.4108 | 0.4003 | 0.4623 | 0.4279 | 0.3979 | 0.3938 | 0.3638 | 0.3838 | 0.4161 | 0.3848 | 0.3744 |
| DNABERT2 | 0.4364 | 0.4617 | 0.4830 | 0.4610 | 0.4606 | 0.4382 | 0.3876 | 0.4387 | 0.4274 | 0.4302 | 0.4352 |
| NT | 0.4524 | 0.4574 | 0.4898 | 0.4741 | 0.4794 | 0.4548 | 0.4091 | 0.4399 | 0.4562 | 0.4312 | 0.4432 |
| Genomics-FM | 0.4724 | 0.4698 | 0.5088 | 0.4867 | 0.4958 | 0.4629 | 0.4227 | 0.4672 | 0.4555 | 0.4552 | 0.4389 |

**Table S19.** Comparison of Genomics-FM and RNA models across various tasks.

| Model | MRL Random Set | MRL Human Set | RNA Switches | Isoform |
| --- | --- | --- | --- | --- |
| Optimus | 0.8400 | 0.7800 | 0.5567 | 0.5082 |
| RNA-FM | 0.8474 | 0.7947 | 0.5598 | 0.7032 |
| Genomics-FM | 0.8498 | 0.7979 | 0.5760 | 0.7319 |

**Table S20.**
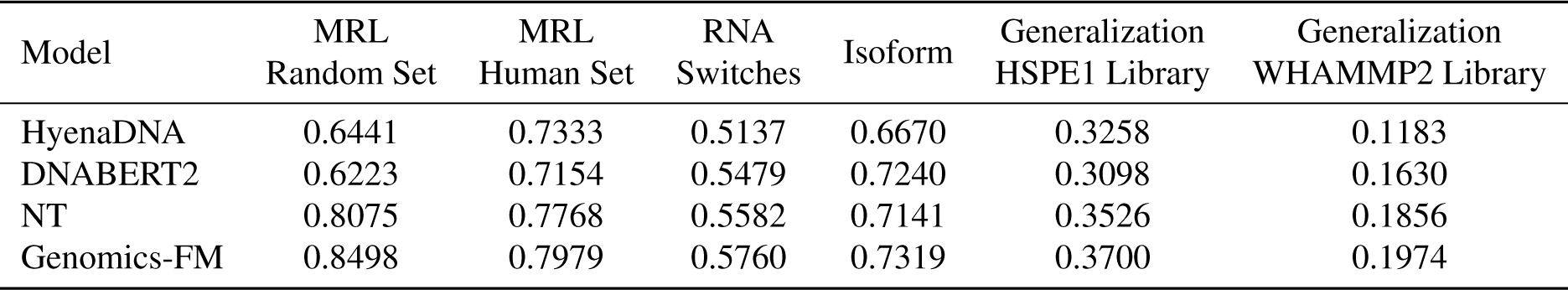
Comparison of Genomics-FM and DNA language models across various tasks.

| Model | MRL Random Set | MRL Human Set | RNA Switches | Isoform | Generalization HSPE1 Library | Generalization WHAMMP2 Library |
| --- | --- | --- | --- | --- | --- | --- |
| HyenaDNA | 0.6441 | 0.7333 | 0.5137 | 0.6670 | 0.3258 | 0.1183 |
| DNABERT2 | 0.6223 | 0.7154 | 0.5479 | 0.7240 | 0.3098 | 0.1630 |
| NT | 0.8075 | 0.7768 | 0.5582 | 0.7141 | 0.3526 | 0.1856 |
| Genomics-FM | 0.8498 | 0.7979 | 0.5760 | 0.7319 | 0.3700 | 0.1974 |

**Table S21.**
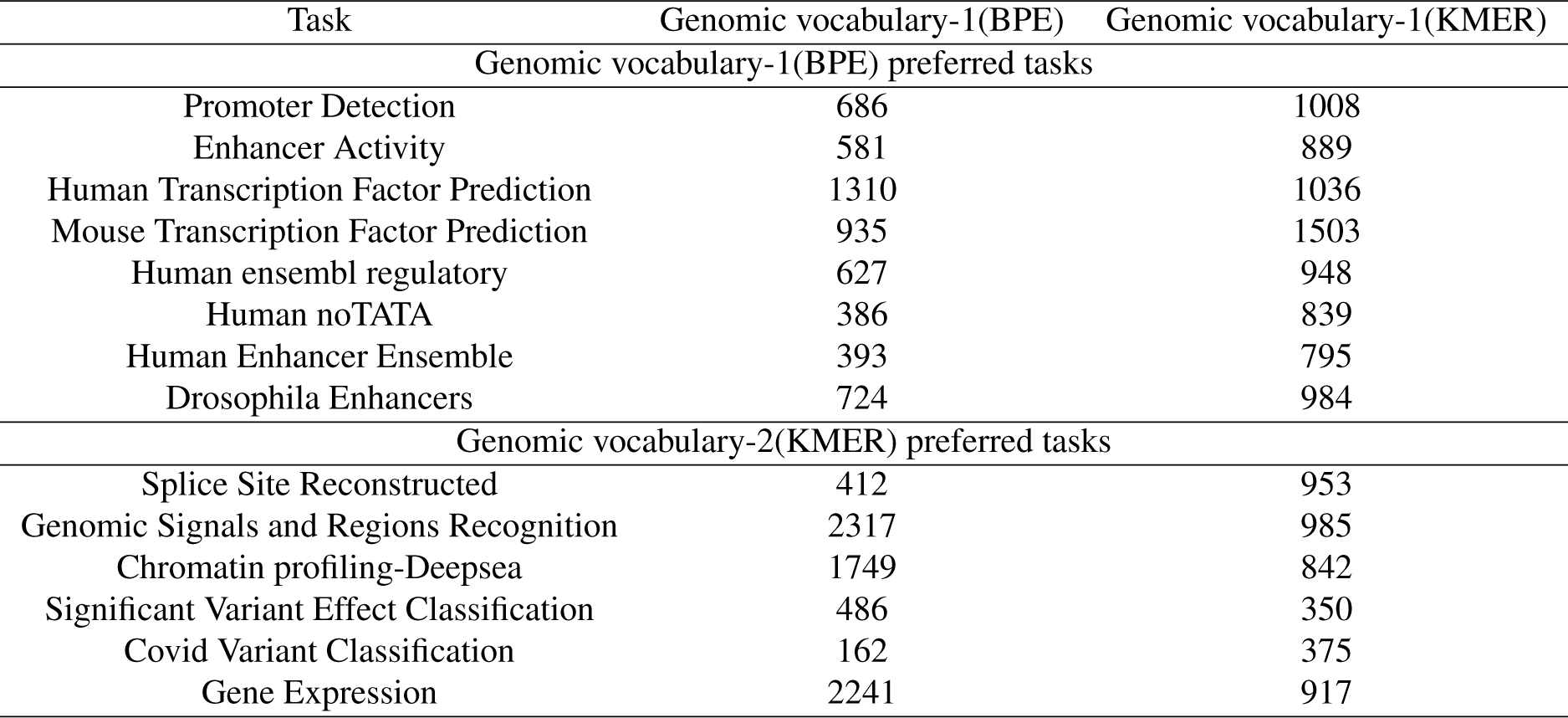
Overlap Between Genomics Vocabularies and Task-Specific Vocabulary in Downstream Tasks.

| Task | Genomic vocabulary-1(BPE) | Genomic vocabulary-1(KMER) |
| --- | --- | --- |
| Genomic vocabulary-1(BPE) preferred tasks |  |  |
| Promoter Detection | 686 | 1008 |
| Enhancer Activity | 581 | 889 |
| Human Transcription Factor Prediction | 1310 | 1036 |
| Mouse Transcription Factor Prediction | 935 | 1503 |
| Human ensembl regulatory | 627 | 948 |
| Human noTATA | 386 | 839 |
| Human Enhancer Ensemble | 393 | 795 |
| Drosophila Enhancers | 724 | 984 |
| Genomic vocabulary-2(KMER) preferred tasks |  |  |
| Splice Site Reconstructed | 412 | 953 |
| Genomic Signals and Regions Recognition | 2317 | 985 |
| Chromatin profiling-Deepsea | 1749 | 842 |
| Significant Variant Effect Classification | 486 | 350 |
| Covid Variant Classification | 162 | 375 |
| Gene Expression | 2241 | 917 |

